# Multi-cohort, multi-sequence harmonisation for cerebrovascular brain age

**DOI:** 10.1101/2025.04.22.649344

**Authors:** Mathijs B.J. Dijsselhof, Candace Moore, Saba Amiri, Mervin Tee, Saima Hilal, Christopher Chen, Bert-Jan H. van den Born, Wibeke Nordhøy, Ole A. Andreassen, Lars T. Westlye, Nishi Chaturvedi, Alun D. Hughes, David M. Cash, Jonathan M. Schott, Carole H. Sudre, Frederik Barkhof, Joost P.A. Kuijer, Francesca Biondo, James H. Cole, Henk J.M.M. Mutsaerts, Jan Petr

## Abstract

**Introduction:** Higher brain-predicted age gaps (BAG), based on anatomical brain scans, have been associated with cognitive decline among elderly participants. Adding a cerebrovascular component, in the form of arterial spin labelling (ASL) perfusion MRI, can improve the BAG predictions and potentially increase sensitivity to cardiovascular health, a contributor to brain ageing and cognitive decline. ASL acquisition differences are likely to influence brain age estimations, and data harmonisation becomes indispensable for multi-cohort brain age studies including ASL. In this multi-cohort, multi-sequence study, we investigate harmonisation methods to improve the generalisability of cerebrovascular brain age.

**Methods:** A multi-study dataset of 2608 participants was used, comprising structural T1-weighted (T1w), FLAIR, and ASL 3T MRI data. The single scanner training dataset consisted of 806 healthy participants, age 50±17, 18-95 years. The testing datasets comprised four cohorts (n=1802, age 67±8, 37-90 years). Image features included grey and white matter (GM/WM) volumes (T1w), WM hyperintensity volumes and counts (FLAIR), and ASL cerebral blood flow (CBF) and its spatial coefficient of variation (sCoV). Feature harmonisation was performed using NeuroComBat, CovBat, NeuroHarmonize, OPNested ComBat, AutoComBat, and RELIEF. ASL-only and T1w+FLAIR+ASL brain age models were trained using ExtraTrees. Model performance was assessed through the mean absolute error (MAE) and mean BAG.

**Results:** ASL feature differences between cohorts decreased after harmonisation for all methods (p<0.05), mostly for RELIEF. Negative associations between age and GM CBF (b=-0.37, R^2^=0.13, unharmonised) increased after harmonisation for all methods (b<-0.42, R^2^>0.12) but weakened for RELIEF (b=-0.28, R^2^=0.14). In the ASL-only model, MAE improved for all harmonisation methods from 11.1±7.5 years to less than 8.8±6.2 years (p<0.001), while BAGs changed from 0.6±13.4 years to less than -1.03±7.92 years (p<0.001).

For T1w+FLAIR+ASL, MAE (5.9±4.6 years, unharmonised) increased for all harmonisation methods non-significantly to above 6.0±4.9 years (p>0.42) and significantly for RELIEF (6.4±5.2 years, p=0.02), while BAGs non-significantly differed from -1.6±7.3 years to between -1.3±4.7 and -2.0±8.0 years (p>0.82). In general, the ASL-specific parameter harmonisation method AutoComBat performed nominally best.

**Discussion:** Harmonisation of ASL features improves feature consistency between studies and also improves brain age estimations when only ASL features are used. ASL-specific parameter harmonisation methods perform nominally better than basic mean and scale adjustment or latent-factor approaches, suggesting that ASL acquisition parameters should be considered when harmonising ASL data. Although multi-modal brain age estimations were improved less by ASL-only harmonisation, possibly due to weaker associations between age and ASL features compared to T1w features feature importance, studies investigating pathological ASL-feature distributions might still benefit from harmonisation. These findings advocate for ASL-parameter specific harmonisation to explore associations between cardiovascular risk factors, brain ageing, and cognitive decline using multi-cohort ASL and cerebrovascular brain age studies.

## 1. Introduction

Ageing is associated with the decline of physiological health and the development of pathology(Franke et al., 2020). The brain appearing older than normal for its chronological age is associated with increased risks of cognitive decline and mortality(Biondo et al., 2022; Cole et al., 2018). The brain-predicted age gap (BAG) is defined as the difference between the neuroimaging-derived predicted biological age and chronological age. BAG has shown value in predicting the risk of neurodegenerative pathology and psychiatric disorders, and determining and monitoring treatment strategies(Baecker et al., 2021). Commonly used neuroimaging-derived features to determine BAG are macroanatomical brain features such as GM and WM volume, and recent approaches have included multi-modality features to increase accuracy and sensitivity to functional and physiological brain ageing, and to predict the development of specific pathologies(Jirsaraie et al., 2023).

Cerebrovascular health markers are good extensions for BAG assessment as cerebrovascular pathology plays a role in many diseases ultimately leading to cognitive decline, such as vascular dementia(Pantoni, 2010) or Alzheimer’s Disease (AD)(Iadecola & Gottesman, 2019; Iturria-Medina et al., 2016). Additionally, it is essential to monitor the effects of cardiovascular health management in the cerebrovascular system(Dolui et al., 2022; Tryambake et al., 2013) due to its direct impact on cerebrovascular health(Williamson et al., 2018). Therefore, incorporating cerebrovascular health features into BAG may increase sensitivity to the risk of cognitive decline and mortality, and provide valuable insights into the impact of cardiovascular health (e.g., blood pressure change, coronary heart disease) and its (interventional) treatment on brain physiology and pathology.

An established method to non-invasively assess cerebrovascular health is Arterial Spin Labeling (ASL) perfusion MRI(Alsop et al., 2015; Clement et al., 2022; Lindner et al., 2023). ASL-derived cerebral blood flow (CBF) has been shown to correlate with cognitive decline(Binnewijzend et al., 2013; van Dinther et al., 2024), amyloid-β and tau pathology(Falcon et al., 2024) and synaptic dysfunction(Falcon et al., 2024). Improved BAG accuracy and classification of AD patients was shown by adding ASL features to commonly used T1w and FLAIR features, dubbed “cerebrovascular brain age”, in a single-cohort study (M. B. J. Dijsselhof et al., 2023; Rokicki et al., 2021). Although promising, these pre-trained single-cohort models do not generalise well to ASL datasets as a large variety of ASL implementations exist, with differences in acquisition hardware, labelling, and readout methods, and acquisition parameters(Grade et al., 2015). Furthermore, ASL acquisition parameters interact with the physiological state(Clement et al., 2018), resulting in additional differences between cohorts. All these factors influence the quantified perfusion values(H. J. M. M. Mutsaerts et al., 2015), introducing undesirable variability to the modelling of the relationship of age or disease with perfusion. One way to circumvent these issues is to train cerebrovascular brain age models using a mixture of datasets that cover the differences in ASL sequence parameters. Unfortunately, individual ASL studies tend to be limited in size, lack healthy controls, and cover a narrow age range(de Lange et al., 2022). Harmonisation is thus needed to reduce site- or sequence-related biases, commonly referred to as batch effects (Hu et al., 2023), and improve the crucial generalisability of cerebrovascular brain age estimations(Gaser et al., 2024).

In recent years, several harmonisation methods were developed that aim to minimise non-biological variance driven by aspects such as acquisition technique difference, while preserving biological and pathological associations and increasing power(Hu et al., 2023). The first methods correct the mean and variance of features across sites, such as NeuroComBat(Fortin et al., 2017), or include between-feature covariance estimates, such as in CovBat(A. A. Chen et al., 2022). Later methods attempted to address and retain complex associations between the harmonised features and biological covariates. NeuroHarmonize(Pomponio et al., 2020) models non-linear correlations with age, addressing possible non-linear correlations of age with arterial transit time (ATT) and CBF. These methods remain limited to a single batch effect estimation, while Complex associations between sites, scanners, and ASL acquisition parameters might need to be resolved through multiple harmonisation steps. OPNested ComBat with multiple ASL-specific batch variables might offer more flexibility to harmonise similar and distinctive ASL sequences(Horng et al., 2022). Furthermore, small differences in ASL acquisition parameters, such as the post-labeling delay (PLD), might result in similar batch effects, and AutoComBat mitigates sequence-dependent variations by clustering subjects into automatically identified batches by assessing several image acquisition parameters (Carré et al., 2022). Lastly, these approaches assume all batch effects are known, while this may not be the case, and identification of latent batch effects might further improve harmonsation. RELIEF incorporates prior batch effect knowledge and estimates latent batch effects(Zhang et al., 2023) to deal with unknown batch effects and mitigate ASL sequence parameter differences. Although some methods have been applied to structural BAG(Lombardi et al., 2020; Marzi et al., 2024; Pomponio et al., 2020), their impact on ASL-related issues with heterogeneity across sites and sequences is still unknown.

Here, we investigate harmonisation methods to improve the generalisability of cerebrovascular brain age in six cohorts differing in age range, and ASL acquisition types and parameters. Specifically, we investigate 1) the ability of harmonisation methods to reduce the between-cohort bias in ASL features, 2) the effect of different harmonisation methods on the accuracy of solely cerebrovascular, and combined structural and cerebrovascular brain age predictions, and 3) similarities in BAG before and after harmonisation.

## 2. Methods

### 2.1 MRI Datasets

Training data were drawn from two cohorts scanned at the same scanner: the healthy controls of the StrokeMRI cohort, obtained at two time points, and the Thematically Organized Psychosis (TOP) cohort(Rokicki et al., 2021). Both studies were approved by the Regional Committee for Medical Research Ethics and the Norwegian Data Inspectorate. Testing data were drawn from several population-based cohorts: the Healthy Life in an Urban Setting (HELIUS)(Snijder et al., 2017); Southall And Brent Revisited (SABRE)(Jones et al., 2020); Epidemiology of Dementia In Singapore (EDIS)(Wong et al., 2020); and Insight 46 (a sub-study of the MRC NSHD; the British 1946 birth cohort)(Lane et al., 2017) studies. The HELIUS study was approved by the Ethical Review Board of the Amsterdam University Medical Center. EDIS was approved by the National Healthcare Group-specific Review Board and the Singapore Eye Research Institute. The National Research Ethics Service (NRES) Committee London granted ethical approval for SABRE (14/LO/0108) and Insight 46 (14/LO/1173). All participants provided written informed consent. Participants with mild cognitive impairment or dementia, as defined per cohort separately, or major brain pathology were excluded.

### 2.2 Imaging acquisition and processing

The study cohorts, MRI scanner and platform, and, sequence parameters of the acquired structural T1-weighted (T1w) and T2-weighted (T2w) Fluid Attenuated Inversion Recovery (FLAIR), and ASL scans in each study can be found in Table 1. Image processing was performed with ExploreASL version 1.11.0 (Mutsaerts et al., 2020) using Statistical Parametric Mapping 12 (SPM12), version r7219. Briefly, tissue segmentation of the structural T1w images into grey matter (GM), white matter (WM), and cerebrospinal fluid (CSF) was performed using the Computational Anatomy Toolbox 12 version r1615(Gaser, 2009). WM hyperintensities (WMH) were segmented from FLAIR and used to fill WMH on T1w images ahead of segmentation using the lesion prediction algorithm of the Lesion Segmentation Toolbox version 2.0.15(Schmidt et al., 2012). WMH volume and count (the number of spatially discrete clusters) were determined. Regions-of-interests (ROI) for GM were created as an intersection of the SPM12 GM, the anterior cerebral artery (ACA), the middle cerebral artery (MCA), and the posterior cerebral artery (PCA)(Tatu et al., 2012) ROIs with the individual CAT12 GM segmentations (partial volume > 0.7). ASL images were rigid-body registered to T1w images. The recommended single-compartment model was used to quantify CBF (Alsop et al., 2015). Partial-volume corrected mean CBF (Asllani et al., 2008) and the ASL sCoV — as the ratio of standard deviation divided by the mean(H. J. Mutsaerts et al., 2017) — were calculated in bilateral total GM and vascular territory ROIs. All images and ROIs were transformed to the Montreal Neurological Institute (MNI) standard space. CBF images were visually checked for typical artefacts(Alsop et al., 2015), such as inefficient labelling, excessive motion, or strong arterial transit time (ATT) artefacts, and if present were excluded from the study. Exclusion examples are shown in Supplementary Figure 1.

**Table 1:**
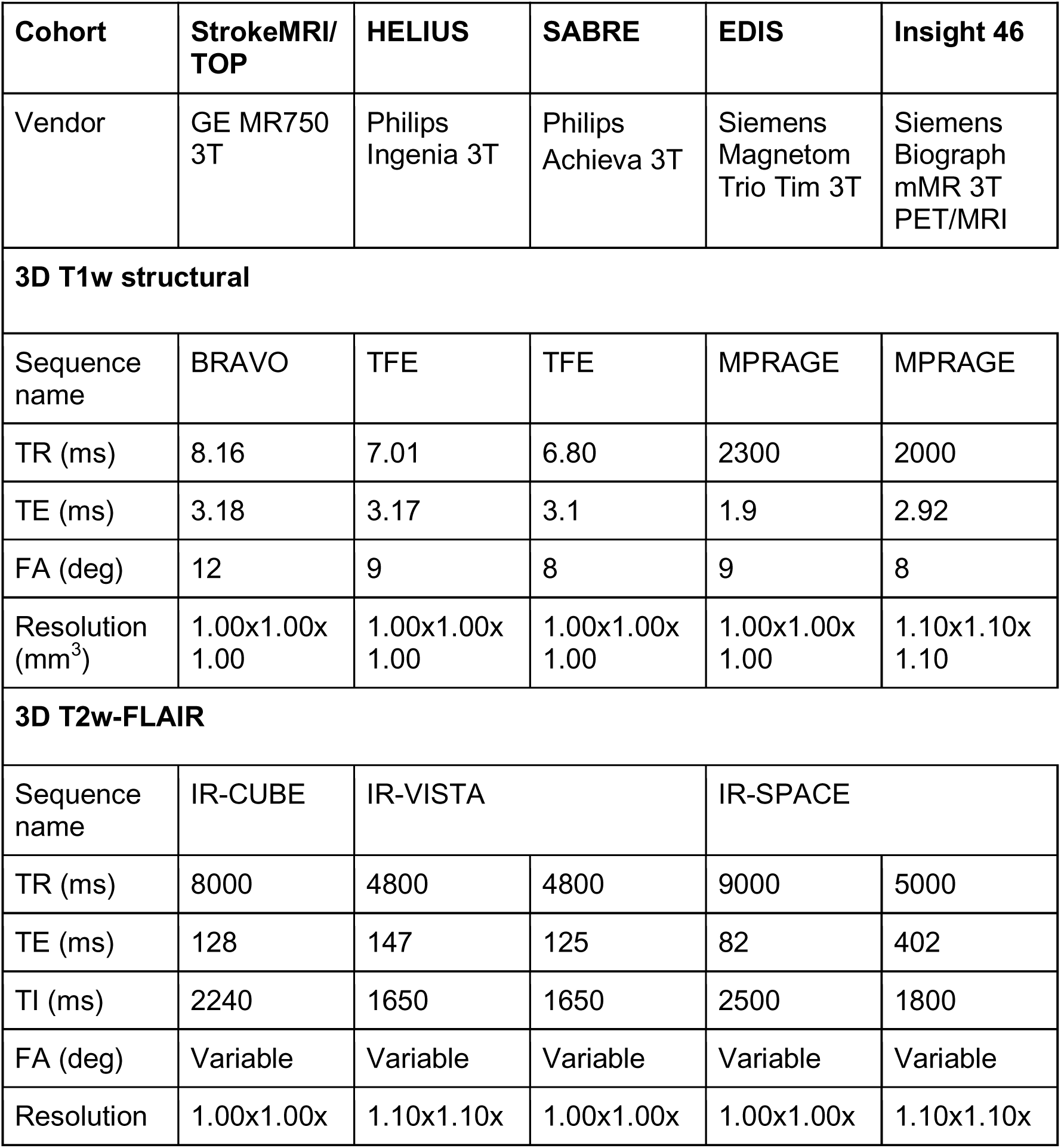
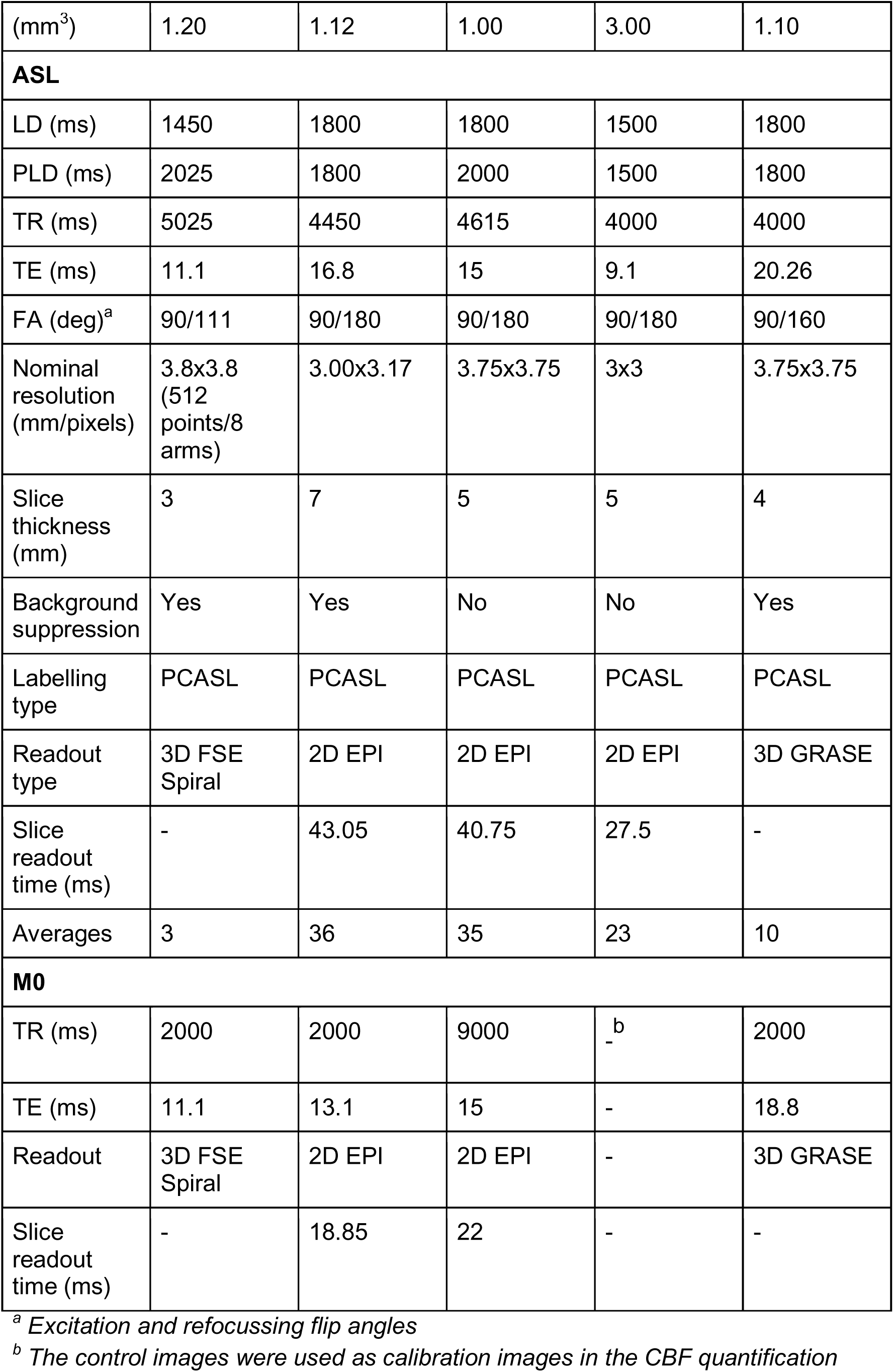
Sequence parameters for the structural—T1w and T2w-FLAIR— and ASL images in the StrokeMRI, TOP, HELIUS, SABRE, EDIS, and Insight 46 cohorts. M0 was not available in the EDIS study and the mean control image was used instead. *ASL: arterial spin labelling; BRAVO: Brain Volume imaging; EPI: echo-planar imaging; FA: flip angle; FSE: fast spin-echo; GRASE: gradient and spin-echo; IR: inversion recovery; LD: labelling delay; MPRAGE: magnetization prepared rapid gradient echo; M0: equilibrium magnetisation image; PCASL: pseudo-continuous ASL; PLD: post-labelling delay; SPGR: spoiled gradient; TE: echo time; TFE: turbo field echo; TI: inversion time; TR: repetition time; T1w: T1-weighted; T2w: T2-weighted*.

### 2.3 Machine learning

To estimate unharmonised BAG, features from ASL-only, or including T1-weighted and FLAIR images together (T1w+FLAIR+ASL) were used. T1w features consisted of GM, WM, and CSF volumes, the ratio of GM to the intracranial volume (ICV), the ratio of both GM and WM to the ICV. Log-transformed FLAIR features consisted of the ratio of WMH volume divided by WM volume, and WMH count. ASL features consisted of both GM and vascular-territory-based CBF and log-transformed sCoV(M. B. J. Dijsselhof et al., 2023).

StrokeMRI and TOP were combined into a single training dataset, as both cohorts were obtained on the same 3T MR750 GE scanner with a 32-channel head coil, with the exact same scanner software release, sequences, and sequence parameters. The training was performed using the ExtraTrees algorithm and the full training dataset, and model performance was estimated through 5-fold cross-validation stratified for age and sex in the same training dataset. The validation results were summarised across all folds and hereafter referred to as the Validation dataset. Testing was performed in the HELIUS, SABRE, EDIS, and Insight 46 cohorts to obtain the BAG and the mean absolute error (MAE; the mean absolute difference between the estimated and chronological brain age), and to determine the coefficient of determination (R^2^). Brain age estimations are commonly biased by the regression-to-the-mean effect(de Lange & Cole, 2020). The age bias was estimated across all cross-validation folds during training by regressing the predicted age on chronological age (Supplementary Figure 2) and subsequently applied to correct the predicted age in all testing datasets(B. Zhang et al., 2023). Training, validation, and testing were performed separately for the ASL-only and T1w+FLAIR+ASL models.

### 2.4 ASL feature harmonisation

We tested the following feature-level statistical harmonisation methods mentioned by He et al.(Hu et al., 2023): NeuroComBat(Fortin et al., 2017), CovBat, (A. A. Chen et al., 2022), NeuroHarmonize(Pomponio et al., 2020), AutoComBat(Carré et al., 2022), OPNested ComBat(Horng et al., 2022), and RELIEF(R. Zhang et al., 2023). The ASL features of all datasets (training and testing) were harmonised for each harmonisation method separately, using age and sex as covariates. After harmonisation, the training (using only the training dataset) and 5-fold cross-validation of the training set were performed for every harmonisation method separately. Next, testing (in EDIS, HELIUS, Insight 46, and SABRE datasets), and the abovementioned age-bias corrections, were repeated per harmonisation method with the corresponding harmonised datasets. This process was repeated for the ASL-only and T1w+FLAIR+ASL models separately. In total, the five datasets (training, EDIS, HELIUS, Insight 46, and SABRE) were harmonised using six methods and brain age was trained for two models (ASL-only or T1w+FLAIR+ASL), resulting in 60 different dataset-harmonisation-model combinations (Figure 1).

**Figure 1:**
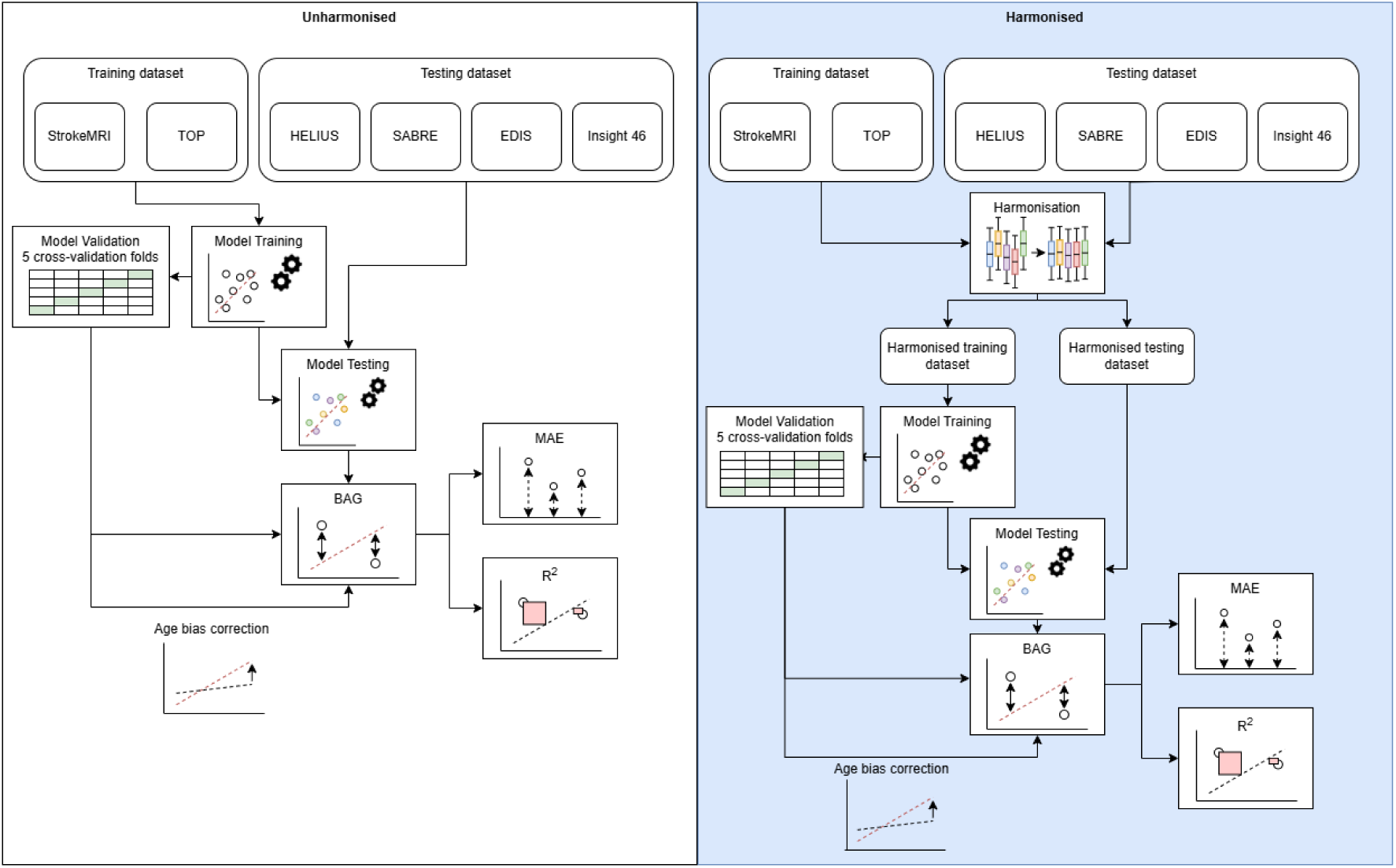
Brain age estimation model training and testing, and model performance evaluation, with (right) and without (left) data harmonisation. *BAG: brain-predicted age gap; MAE: mean absolute error*.

### 2.5 Statistical analyses

#### 2.5.1 Demographics

Statistical analyses were conducted in R version 4.3.1 (R Core Team, 2023). Before harmonisation, all imaging data were tested for normal distribution using the Shapiro-Wilk test. The ratio of WMH volume to WM volume, WMH count, and all sCoV features were log-transformed because of their right-skewed distributions. To test if T1w, FLAIR, and ASL features differed between all cohorts before harmonisation, analysis of covariance (ANCOVA) corrected for age and sex with Tukey post-hoc tests corrected were performed.

#### 2.5.2. ASL feature harmonisation

To compare the overall effect of each harmonisation method, the mean of ASL features across all cohorts were compared between harmonisation methods using ANCOVA corrected for age and sex with Tukey post-hoc tests. To compare the effect of each harmonisation method on the difference of ASL features between cohorts, ANCOVA corrected for age and sex with Tukey post-hoc tests for all cohort combinations (cohort-pairs) were performed separately for each ASL feature and harmonisation method. To investigate the association between age and change in each ASL feature between unharmonised and harmonisation methods, linear regressions were performed. Additionally, an interaction term between each ASL feature and cohort was used to investigate differences in the association between the training dataset and each testing dataset.

#### 2.5.3. Brain age

To assess the effect of harmonisation on the ASL-only model performance, differences of MAE in the validation dataset were compared between every harmonisation method and without harmonisation, using ANOVA with Tukey post-hoc tests. Similar tests were performed to assess the performance in the testing dataset.

The following analyses were performed in the testing datasets only. To investigate if harmonisation changed the explained variance within each model across all cohorts, the R^2^ was determined. The effect of each harmonisation method on the MAE difference between cohorts was investigated using ANOVA with Tukey post-hoc tests for all cohort combinations (cohort-pairs). The same methods were used to assess the effect of each harmonisation method on the difference of BAG between unharmonised and harmonised data, or difference of BAG between cohorts for each harmonisation method, or unharmonised separately. These analyses have been performed for the models using ASL-only and T1w+FLAIR+ASL features separately.

### 2.6. Sensitivity analyses

To understand the effect of age bias correction, all brain age statistical analyses comparing unharmonised and harmonised data across all cohorts were repeated for BAG and MAE values not corrected for age bias.

### 2.7. Post-hoc

The effect of ASL-feature harmonisation on the overall MAE and BAG in the T1w+FLAIR+ASL models might be limited due to possible high importance of the structural features in the model. In this case, a post-hoc analysis will be performed to investigate the underlying reasons. The importance of the structural features in determining the BAG, which will be assessed using Shapley values(Rozemberczki et al., 2022). To explore this hypothesis, the effect of NeuroComBat harmonisation, as the most commonly used harmonisation method(Hu et al., 2023), on all T1w, FLAIR, and ASL features will be investigated additionally. T1w+FLAIR+ASL model-derived MAE and BAG will be compared between unharmonised and NeuroComBat harmonised features using t-tests, and the R^2^ calculated. Differences in MAE and BAG between cohorts for the NeuroComBat harmonised results will be compared using ANOVA with Tukey post-hoc tests. In all statistics, p<0.05 was defined as statistically significant.

## 3. Results

### 3.1 Demographics

After the exclusion of 974 participants (Training: 17; HELIUS: 40; SABRE: 86; EDIS: 611; Insight 46: 220; Supplementary Figure 1), a total of 2608 participants remained (Figure 2, Table 2). All training dataset features differed between the population datasets (p < 0.05) before harmonisation, except for the WMH count between Training and HELIUS datasets (p = 0.80), ACA CBF between the Training and EDIS (p = 0.49), and PCA CBF between training and Insight 46 (p = 0.15). Cohort differences can be found in Supplementary Table 1.

**Figure 2:**
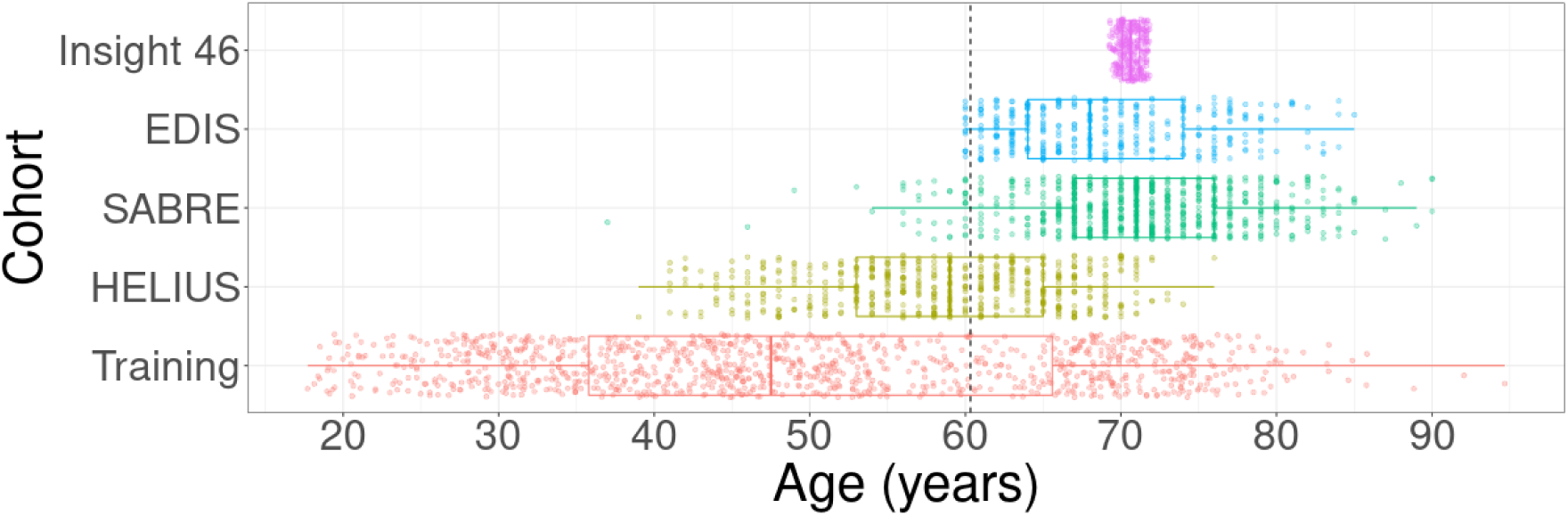
Age distribution per cohort. The training dataset consists of StrokeMRI and TOP combined. The dotted line represents the mean age of all cohorts combined.

**Table 2:**
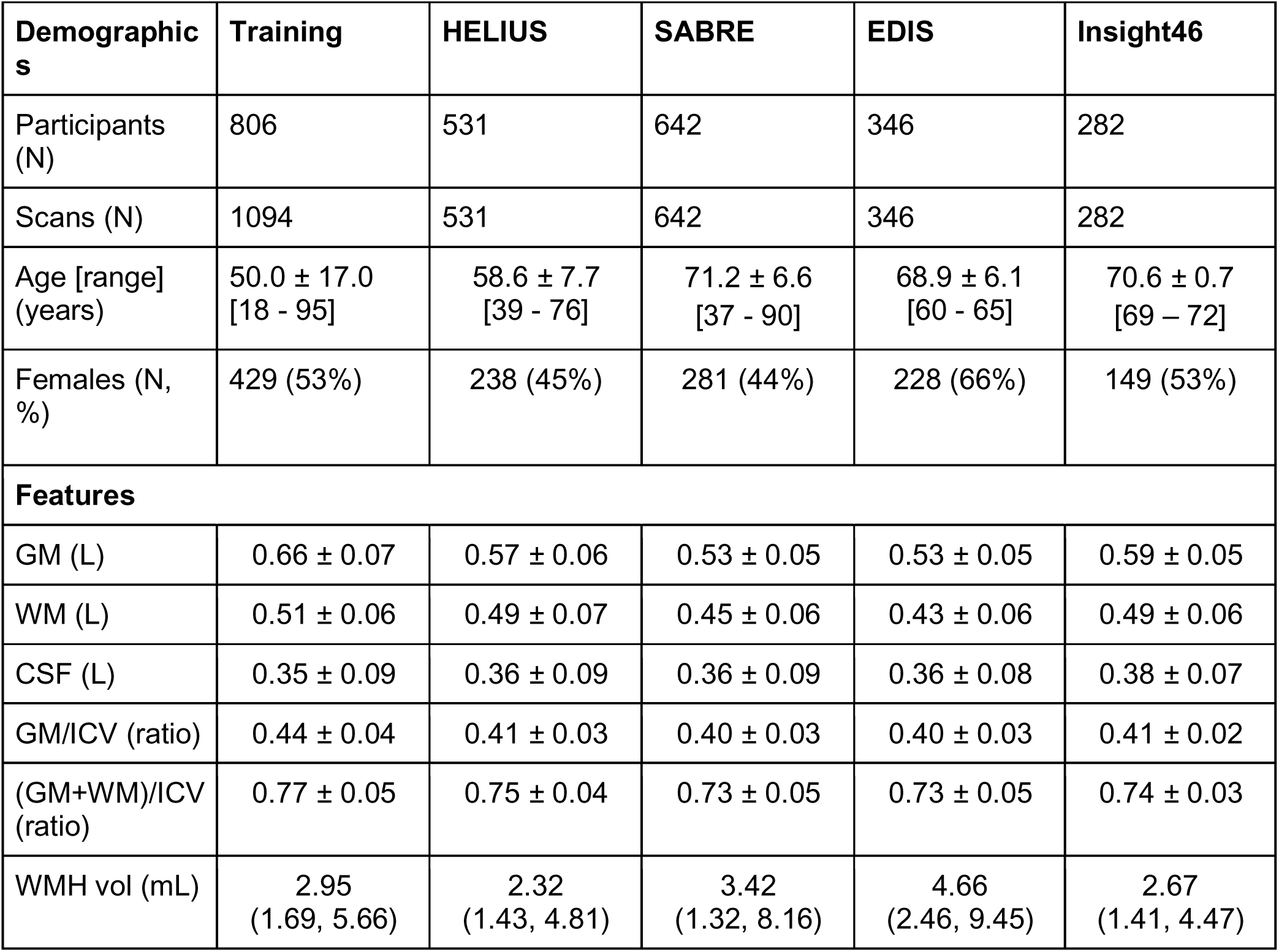
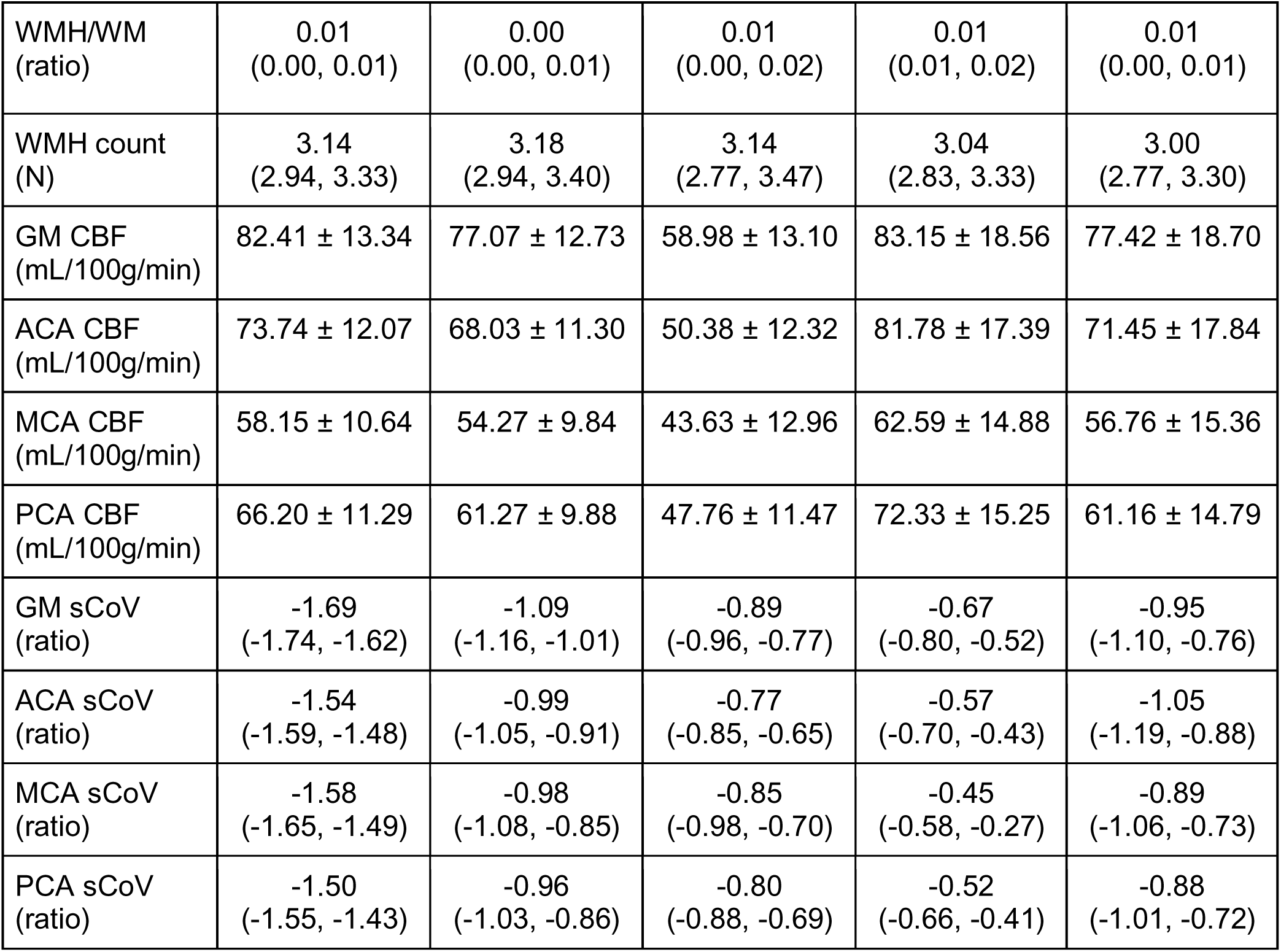
Demographics and imaging derivatives of the training and testing datasets. Unless stated otherwise, data are mean ± standard deviations or median (interquartile range). WMH/WM, WMH count, and all sCoV features have been log-transformed. *ACA: anterior cerebral artery; CBF: cerebral blood flow; CSF: cerebrospinal fluid; GM: grey matter; ICV: intracranial volume; MCA: middle cerebral artery; PCA: posterior cerebral artery; sCoV: spatial coefficient of variation; WM: white matter; WMH: white matter hyperintensities*.

### 3.2 Feature harmonisation

For all cohorts combined (i.e. the full testing dataset), no differences in mean CBF and sCoV values (p > 0.05) were found between each harmonisation method and without harmonisation, however, the distribution was reduced after harmonisation (Supplementary Table 2A).

GM CBF was different between 9 out of 10 cohort-pairs (p < 0.001) before harmonisation. The difference between cohorts in GM CBF decreased after harmonisation with 7 cohort-pairs being different for NeuroCombat, CovBat, NeuroHarmonize, and AutoComBat (p < 0.001); 6 cohort-pairs for OPNested ComBat (p < 0.001), and no cohort-pairs for RELIEF (p > 0.99) (Figure 3). The same results applied to ACA, MCA, and PCA CBF, with the exceptions of ACA CBF being different between 8 cohort-pairs (p < 0.001) before harmonisation; 6 cohort-pairs for NeuroHarmonize (p < 0.01), and PCA CBF being different 5 cohort-pairs for OPNested ComBat (p < 0.01). All cohort and harmonisation-specific CBF values can be found in the Supplementary Table 2B.

**Figure 3:**
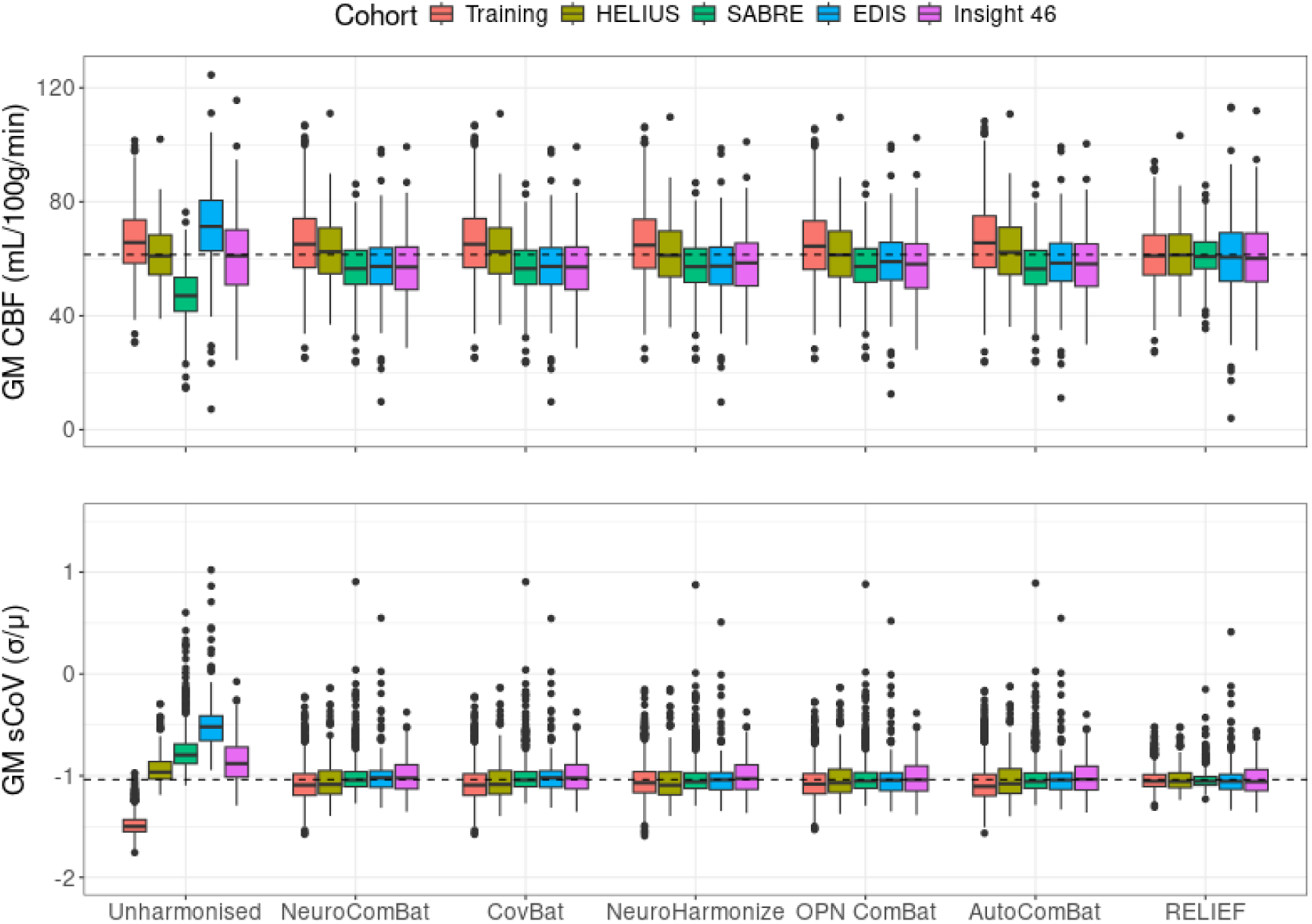
Boxplots of GM CBF and sCoV per cohort for every harmonisation technique. The box describes the second and third quartile range with the median is The dashed line represents the average unharmonised CBF value of all cohorts combined. *CBF: cerebral blood flow; GM: grey matter; sCoV: spatial coefficient of variation*.

GM sCoV was different between all cohorts (p < 0.001) before harmonisation. The difference between cohorts in GM sCoV decreased after harmonisation (Supplementary Table 2A): with 6 cohort-pairs being different NeuroComBat and CovBat (p < 0.01); 4 cohort-pairs for NeuroHarmonize, OPNested ComBat, and AutoComBat (p < 0.03); and, no cohort-pairs for RELIEF (p > 0.99) (Figure 3). The same results applied to ACA, MCA, and PCA sCoV, with the exceptions of ACA sCoV being different between 5 cohort-pairs for NeuroComBat and CovBat (p < 0.04); 2 cohort-pairs for NeuroHarmonize, AutoComBat, and OPNested ComBat (p < 0.04); MCA sCoV being different between 3 cohort-pairs for NeuroComBat and CovBat (p < 0.05); 1 cohort-pair for OPNested ComBat (p < 0.02), and between no cohorts for NeuroHarmonize and AutoComBat (p > 0.16). Lastly, PCA sCoV was different between 7 cohort-pairs for NeuroComBat and CovBat (p < 0.05); 5 cohort-pairs for OPNested ComBat (p < 0.02), and 6 cohort-pairs for AutoComBat (p < 0.04). All cohort and harmonisation-specific sCoV values can be found in the Supplementary Table 2B.

Associations between age and GM CBF changed before and after harmonizations (Figure 4A, Supplementary Table 3A). Age was associated with GM CBF (b = -0.37, R^2^ = 0.13, p < 0.001) before harmonisation, and the association nominally increased for NeuroComBat, CovBat, NeuroHarmonize, and OPNested ComBat (b between - 0.42 and -0.45, R^2^ between 0.12 and 0.15, p < 0.001) and even more for AutoComBat (b = -0.47, R^2^ = 0.17, p < 0.001), but decreased for RELIEF (b = -0.28, R^2^ = 0.14, p < 0.001). Age was similarly associated (p < 0.001) with ACA and MCA CBF (b = -0.31, R^2^ = 0.13), and PCA CBF (b = -0.29, R^2^ = 0.07) before harmonisation. Similar behavior of the harmonisation methods to GM CBF were observed for MCA and PCA CBF. For ACA CBF, the association with age decreased for NeuroCombat, CovBat and NeuroHarmonize, OPNested ComBat, and RELIEF, (b between -0.17 and -0.30, R^2^ between 0.02 and 0.09) while increasing for AutoComBat (b = -0.32, R^2^ = 0.10). The associations between age and the CBF features differed (p < 0.001) between the training and testing datasets for all harmonisation methods.

**Figure 4:**
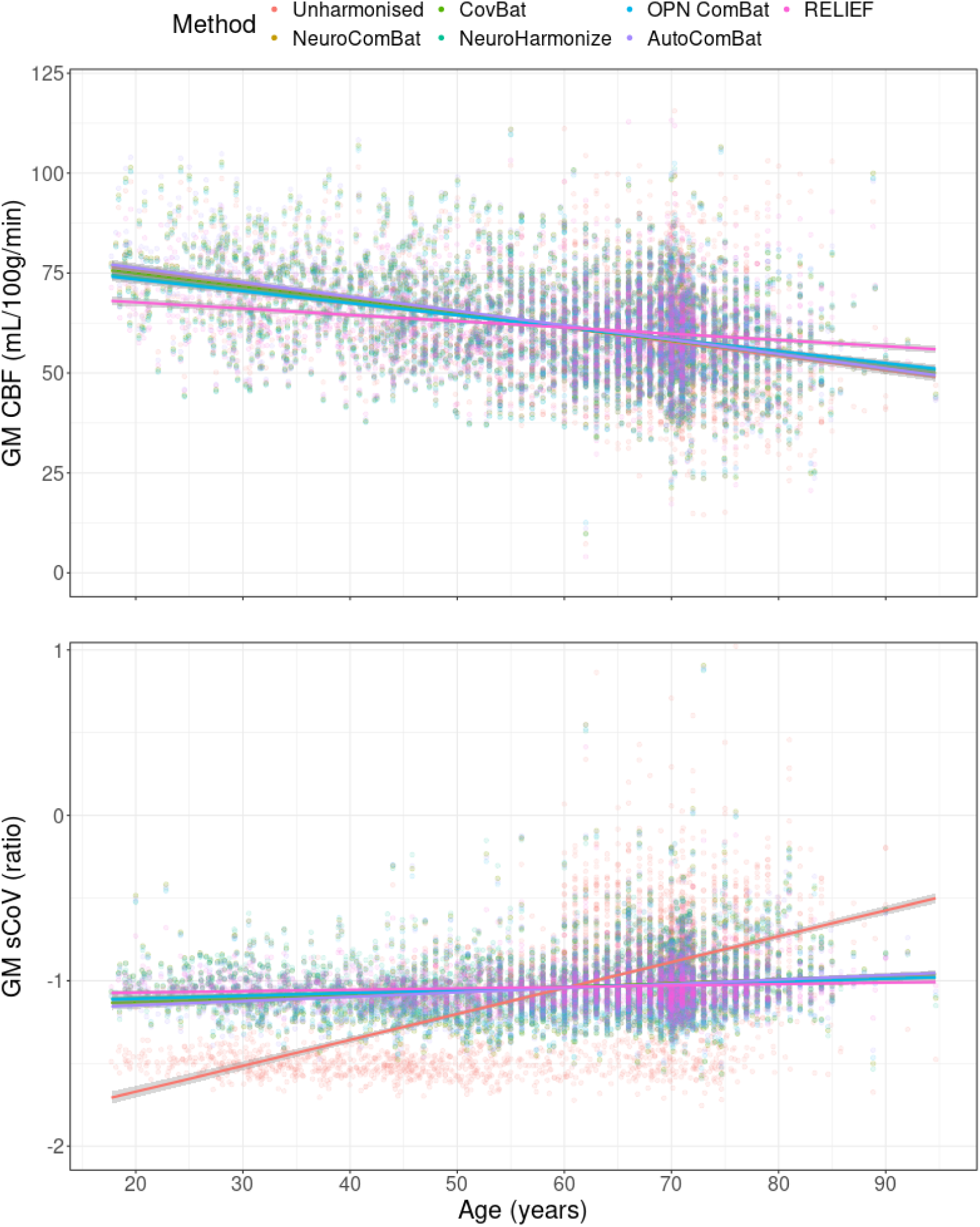
The associations of age with CBF and log-transformed sCoV in GM within the full testing dataset (n=1801). *CBF: cerebral blood flow; GM: grey matter; sCoV: spatial coefficient of variation*.

Associations between age and sCoV values also changed before and after harmonisations (Figure 4B, Supplementary Table 3B). Age was associated with GM sCoV (b = 20.53, R^2^ = 0.32, p < 0.001) before harmonisation, and the association decreased for all methods (b between 10.8 and 16.0, R^2^ between 0.01 and 0.04) with AutoComBat retaining the strongest association (b = 16.0, R^2^ = 0.04). Age was similarly associated (p < 0.001) with ACA (b = 18.8, R^2^ = 0.32), MCA (b = 18.9, R^2^ = 0.29), and PCA sCoV (b = 18.0, R^2^ = 0.32) before harmonisation. Similar behavior of the harmonisation methods to GM sCoV were observed for ACA, MCA, and PCA sCoV, except NeuroComBat retained the strongest association with ACA sCoV (b = 8.7, R^2^ = 0.01) and RELIEF with MCA sCoV (b = 5.7, R^2^ = 0.01)

The associations between age and GM sCoV differed between the training, EDIS and Insight 46 for NeuroHarmonize (p <0.05), and between all datasets for AutoComBat and RELIEF (p < 0.01). The associations of age with ACA differed the least between cohorts, followed by MCA and PCA after harmonisation (data not shown).

### 3.3 Brain age

#### 3.3.1 ASL-only model

Using only ASL features (CBF and sCoV), the MAE (Table 3A, Figure 5A) of unharmonised data was 10.65 ± 7.50 years in the validation set, and 11.08 ± 7.47 years in the testing set. In the validation dataset, the MAE did not differ statistically after harmonisation for all methods (p > 0.37, compared with unharmonised data). In the testing dataset, MAE differed statistically after harmonisation for all methods (p < 0.001, compared with unharmonised data), with 6.39 ± 4.87 years for NeuroComBat, 6.42 ± 4.87 years for CovBat, 6.32 ± 4.78 years for NeuroHarmonize, 6.54 ± 5.14 years for OPNested ComBat, 6.31 ± 4.89 years for AutoCombat, and 8.78 ± 6.15 years for RELIEF. RELIEF harmonisation differed statistically (p < 0.001) from all other methods, and all the other methods did not differ amongst themselves (p < 0.87, Supplementary Table 4A).

**Figure 5:**
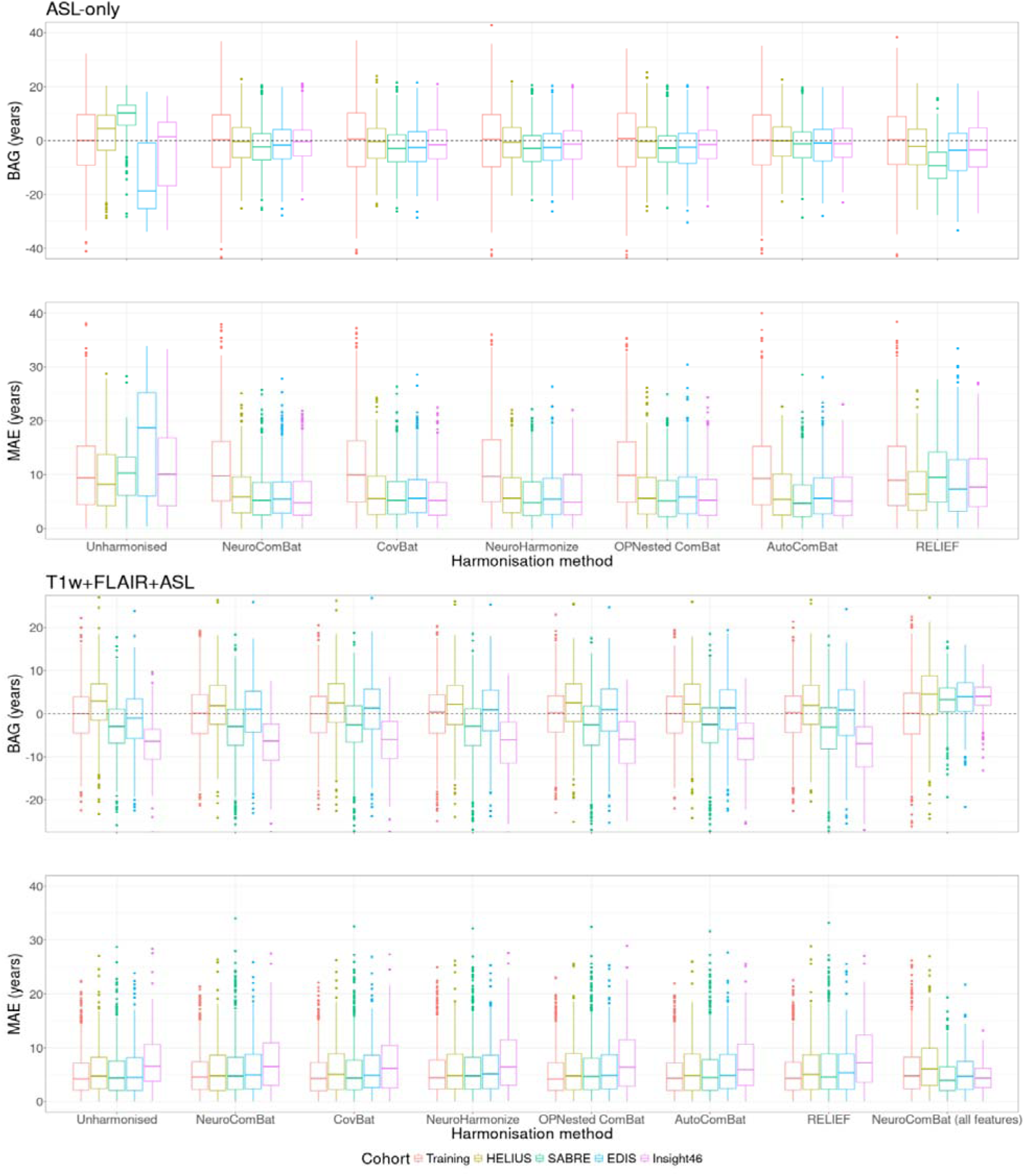
Boxplots of BAG and MAE of all cohorts per harmonisation method, obtained using ASL-only or T1w+FLAIR+ASL features to predict brain age. Harmonisation was performed on ASL features only. Additional results of harmonisation of all features using NeuroComBat have been included under ‘NeuroComBat (all features)’. Note that the validation set was not corrected for age-bias. *ASL: arterial spin labelling; BAG: brain-predicted age gap; MAE: mean absolute error*.

**Table 3:** BAG and MAE per harmonisation method for validation and testing sets, obtained using ASL-only or T1w+FLAIR+ASL features to predict brain age. Additional results of harmonisation of all features using NeuroComBat have been included under ‘NeuroComBat (all features)’. Note that the validation set was not corrected for age-bias. *ASL: arterial spin labelling; BAG: brain-predicted age gap; FLAIR: fluid attenuated inversion recovery; MAE: mean absolute error; T1w: T1-weighted*.

| **ASL-only** | **BAG**<br>**($\mu \pm \sigma$)** | **MAE**<br>**($\mu \pm \sigma$)** | **R<sup>2</sup>** |
| --- | --- | --- | --- |
| **Validation dataset** |  |  |  |
| Unharmonised | -0.23 ± 13.03 | 10.65 ± 7.50 | 0.40 |
| NeuroComBat | -0.43 ± 13.88 | 11.32 ± 8.04 | 0.31 |
| CovBat | -0.22 ± 13.74 | 11.19 ± 7.96 | 0.33 |
| NeuroHarmonize | -0.24 ± 13.90 | 11.37 ± 8.00 | 0.31 |
| OPNested ComBat | -0.12 ± 13.62 | 11.15 ± 7.81 | 0.34 |
| AutoComBat | -0.18 ± 13.06 | 10.62 ± 7.61 | 0.39 |
| RELIEF | -0.22 ± 12.91 | 10.42 ± 7.62 | 0.41 |
| **Testing dataset** |  |  |  |
| Unharmonised | 0.61 ± 13.35 | 11.08 ± 7.47 | 0.23 |
| NeuroComBat | -1.52 ± 7.89 | 6.39 ± 4.87 | 0.48 |
| CovBat | -1.87 ± 7.84 | 6.42 ± 4.87 | 0.48 |
| NeuroHarmonize | -1.76 ± 7.73 | 6.32 ± 4.78 | 0.47 |
| OPNested ComBat | -1.94 ± 8.09 | 6.54 ± 5.14 | 0.44 |
| AutoComBat | -1.03 ± 7.92 | 6.31 ± 4.89 | 0.47 |
| RELIEF | -5.12 ± 9.42 | 8.78 ± 6.15 | 0.33 |
| **T1w+FLAIR+ASL** | **BAG**<br>**($\mu \pm \sigma$)** | **MAE**<br>**($\mu \pm \sigma$)** | **R<sup>2</sup>** |
| **Validation dataset** |  |  |  |
| Unharmonised | -0.07 ± 6.47 | 5.11 ± 3.98 | 0.85 |
| NeuroComBat | 0.11 ± 6.69 | 5.33 ± 4.03 | 0.84 |
| CovBat | -0.08 ± 6.52 | 5.13 ± 4.01 | 0.85 |
| NeuroHarmonize | 0.11 ± 6.78 | 5.39 ± 4.11 | 0.84 |
| OPNested ComBat | $-0.06 \pm 6.56$ | $5.15 \pm 4.06$ | 0.85 |
| AutoComBat | $-0.05 \pm 6.5$ | $5.12 \pm 4.00$ | 0.85 |
| RELIEF | $-0.05 \pm 6.61$ | $5.21 \pm 4.06$ | 0.85 |
| NeuroComBat (all features) | $0.00 \pm 7.30$ | $5.77 \pm 4.47$ | 0.81 |
| **Testing dataset** |  |  |  |
| Unharmonised | $-1.63 \pm 7.28$ | $5.88 \pm 4.60$ | 0.43 |
| NeuroComBat | $-1.75 \pm 7.75$ | $6.18 \pm 4.99$ | 0.43 |
| CovBat | $-1.30 \pm 7.67$ | $6.04 \pm 4.91$ | 0.43 |
| NeuroHarmonize | $-1.70 \pm 7.78$ | $6.21 \pm 4.98$ | 0.43 |
| OPNested ComBat | $-1.50 \pm 7.76$ | $6.15 \pm 4.97$ | 0.42 |
| AutoComBat | $-1.36 \pm 7.64$ | $6.04 \pm 4.88$ | 0.43 |
| RELIEF | $-2.00 \pm 8.01$ | $6.42 \pm 5.17$ | 0.41 |
| NeuroComBat (all features) | $3.56 \pm 5.51$ | $5.31 \pm 3.85$ | 0.63 |

In the testing dataset before harmonisation, the MAE of 5 out of 6 cohort-pairs were statistically different (p < 0.01, Supplementary Table 5A). After harmonisation, the MAE of no cohort-pairs were statistically different for NeuroComBat, CovBat, NeuroHarmonize, and OPNested ComBat (p > 0.13), 2 cohort-pairs for AutoComBat (p < 0.03) and 5 cohort-pairs for RELIEF (p = 0.99). In the validation set, R^2^ did not change for RELIEF, however decreased nominally for all other harmonisation methods compared with unharmonised data. In the testing set, R^2^ improved nominally for all harmonisation methods (Table 3A).

The unharmonised BAGs were -0.23 ± 13.03 and 0.61 ± 13.35 years for the validation and testing sets, respectively (Table 3A, Figure 5A). In the validation dataset, BAGs did not differ statistically after harmonisation for all methods (p > 0.99, compared with unharmonised data). In the testing dataset, BAGs differed statistically after harmonisation for all methods (p < 0.001, compared with unharmonised data), with -1.52 ± 7.89 years for NeuroComBat, -1.87 ± 7.84 years for CovBat, -1.76 ± 7.73 years for NeuroHarmonize, -1.94 ± 8.09 years for OPNested ComBat, -1.03 ± 7.92 years for AutoComBat, and -5.12 ± 9.42 years for RELIEF. No harmonisation methods differed, except for RELIEF, which differed statistically from all other methods (p < 0.001) and AutoComBat, which differed from OPNested ComBat (p = 0.04, Supplementary Table 4A).

In the testing dataset before harmonisation, BAGs of all cohort-pairs were different (p < 0.001, Supplementary Table 5A). After harmonisation, BAGs of 2 cohort-pairs differed for NeuroComBat and CovBat (p < 0.05), 3 cohort-pairs for NeuroHarmonize (p < 0.01), and 4 cohort-pairs for OPNested ComBat (p < 0.03) and RELIEF (p < 0.01). No cohort-pairs were different for AutoComBat (p > 0.06).

#### 3.3.2 T1+ASL+FLAIR model

Using all features (T1w, FLAIR, CBF and sCoV), the MAE (Table 3B, Figure 5B) of unharmonised data was 5.11 ± 3.98 years in the validation set, and 5.88 ± 4.60 years in the testing set. In the validation dataset, the MAE did not differ statistically after harmonisation for all methods (p > 0.71, compared with unharmonised data). In the testing dataset, the MAE was not statistically different (p > 0.37) with 6.18 ± 4.99 years for NeuroComBat, 6.04 ± 4.91 years for CovBat, 6.21 ± 4.98 years for NeuroHarmonize, 6.15 ± 4.97 years for OPNested ComBat, 6.04 ± 4.88 years for AutoComBat, and increased to 6.42 ± 5.17 years for RELIEF (p = 0.02) compared with the unharmonised data. No harmonisation methods were statistically different (p > 0.23, Supplementary Table 4B). In the testing dataset before harmonisation, the MAE of 3 cohort-pairs were different (p < 0.01, Supplementary Table 5B), and remained different after harmonisation (p < 0.03). In the validation set, R^2^ did not differ for all methods except for OPNested ComBat and RELIEF, where it nominally decreased, compared with unharmonised data (Table 3B). Similar results were obtained in the testing set.

The unharmonised BAGs (Table 3B, Figure 5B) were respectively -0.07 ± 6.47 years and -1.63 ± 7.28 years for the validation and testing sets. In the validation dataset, BAGs did not differ statistically after harmonisation for all methods (p > 0.99, compared with unharmonised data). In the testing dataset, BAGs did not differ statistically after harmonisation for all methods (p > 0.78, compared with unharmonised data), with -1.75 ± 7.75 years for NeuroComBat, -1.30 ± 7.67 years for CovBat, -1.70 ± 7.778years for NeuroHarmonize, -1.50 ± 7.76 years for OPNested ComBat, -1.36 ± 7.64 years for AutoComBat, and -2.00 ± 8.01 years for RELIEF. No BAGs of any harmonisation methods were different (p > 0.16, Supplementary Table 4B).

In the testing dataset before harmonisation, BAGs of all cohort-pairs were different (p < 0.001, Supplementary Table 5B), and remained different after harmonisation (p < 0.01).

#### 3.3.3 Sensitivity

Without age-bias correction, in the unharmonised testing dataset of the ASL-only model, the MAE was 13.24 ± 12.28 years (R^2^ = 0.02) and the BAGs were -10.46 ± 14.72 years. After harmonisation, the MAE increased to above 14.66 ± 8.68 years for NeuroComBat, CovBat, NeuroHarmonize, OPNested ComBat, and RELIEF (p < 0.001), while AutoComBat did not differ (13.31 ± 8.17 years, p = 0.99, Supplementary Table 6A). The BAGs decreased to below 12.02 ± 9.96 years (p < 0.001) for all methods. The R^2^ increased to between 0.03 and 0.08 for all methods except for RELIEF, which remained at 0.02.

In the unharmonised testing dataset of the T1w+ASL+FLAIR model, the MAE was 7.4 ± 5.42 years (R^2^ = 0.30), and the BAGs were -4.7 ± 7.88 years. After harmonisation, the MAE and BAGs did not differ for any method (p > 0.61, Supplementary Table 6B). The R^2^ increased to between 0.31 and 0.32 for all methods except RELIEF, which remained at 0.30.

#### 3.3.4 Post-hoc analyses

The three most important features in the T1w+FLAIR+ASL model were the ratio of GM with ICV, the ratio of GM and WM combined with ICV, and the ratio of WMH volume with WM volume (Supplementary Figure 3). After harmonisation of all features (T1w, FLAIR, and ASL) using NeuroComBat, the MAE (Table 3B, Figure 5B) increased to 5.77 ± 4.47 years in the validation set (p < 0.01, compared with unharmonised data). In the testing dataset, the MAE decreased to 5.31 ± 3.85 years (p < 0.01) and differed statistically between 4 out of 6 cohort-pairs (p < 0.03). The BAGs did not change in the validation dataset with 0.00 ± 7.30 years (p = 0.99, compared with unharmonised data). In the testing dataset, the BAGs increased to 3.56 ± 5.51 years (p < 0.001) and differed statistically for 1 out of 6 cohort-pairs (p < 0.01). R^2^ decreased in the validation set and increased in the testing set compared with unharmonised data (Table 3B).

## 4. Discussion

### 4.1 Summary of results

We have investigated six feature-harmonisation methods and their effect on the generalisability of cerebrovascular brain age across five datasets differing in age distribution and ASL acquisition parameters. This study has three main findings. First, all harmonisation methods decreased the difference in ASL features (CBF and sCoV). RELIEF reduced differences in ASL features between cohorts the most. However, it also reduced the association between age and ASL features while all other harmonisation methods retained or even strengthened these associations. Second, cerebrovascular brain age performance improved (lower MAE) in all testing datasets for the ASL-only model after harmonisation. However, this was not the case for the combined (T1w+FLAIR+ASL) cerebrovascular brain age model, while harmonising all features harmonising all features did improve the cerebrovascular brain age performance. Third, with respect to the unharmonised counterpart, BAGs decreased for all methods in the ASL-only model and did not change in the combined model after harmonisation of only ASL features, but increased in the combined model after harmonisation of all features.

### 4.2 Features

Although harmonisation methods reduced mean cohort differences and standard deviations in ASL features, RELIEF harmonisation consistently achieved lower cohort differences. One explanation could be that inherent age-related CBF and sCoV differences between the cohorts are removed instead of preserved in RELIEF as its latent-factor approach does not force its latent variations to be independent of covariates, even when covariates such as age and sex are provided(R. Zhang et al., 2023). It is known that CBF decreases with age(J. J. Chen et al., 2011), and this pattern remains with all harmonisation methods except RELIEF. Moreover, RELIEF reduced associations between age and ASL features, whereas OPNested ComBat and AutoComBat showed the strongest age associations. The likely explanation is that OPNested ComBat and AutoComBat additionally accounted for the ASL acquisition parameters differences between cohorts, which are known to strongly affect their CBF measurements(H. J. M. M. Mutsaerts et al., 2015). The PLD plays a critical role in CBF quantification accuracy, especially in older and cerebrovascularly diseased populations, to account for arterial transit time prolongation that otherwise would result in underestimated CBF(Alsop et al., 2015).

The highest harmonisation effect on CBF and sCoV was found in the SABRE and EDIS cohorts. Unlike the training and other testing datasets with 3D readout, CBF in these datasets was acquired using a 2D EPI readout. This aligns with previously observed CBF measurement differences between 2D and 3D sequences, even when acquired on the same scanner(Baas et al., 2021). Because we used large regions, we expect this to be due to PLD differences between acquired slices or background suppression efficiency (effect of head motion) and M0 estimation rather than the effective spatial resolution and geometric distortion differences between 2D and 3D ASL acquisitions(H. J. M. M. Mutsaerts et al., 2014). The EDIS dataset also had considerably shorter PLD, which, combined with the lack of background suppression, could result in more noise, perhaps explaining the large harmonisation effect. Interestingly, the HELIUS dataset, scanned with a 2D EPI readout including background suppression, was less affected by harmonisation, likely due to the fact that its age range and PLD were more similar to that of the training dataset. AutoComBat, a harmonisation method that utilises ASL parameters to estimate batch effects obtained nominally the best results. This method may be especially useful in older populations, in the presence of pathology, and in 2D ASL acquisitions where the effects of PLD on measured CBF due to increased arterial transit time tend to be stronger(Damestani et al., 2023).

### 4.3 Brain age

Consistent with previous studies, brain age estimations using ASL-only features showed relatively low performance (higher MAE) compared with models using T1w, FLAIR, and ASL features(M. B. J. Dijsselhof et al., 2023; Rokicki et al., 2021), which can be attributed to the high physiological variability of perfusion(Clement et al., 2018). In the ASL-only model, validation dataset performance was not different after harmonisation although its variance was increased. As the training dataset had more stringent criteria for selecting healthy participants than the other included datasets(Rokicki et al., 2021), patterns of general ageing from the testing datasets were possibly introduced to the training dataset through the joint harmonisation of all datasets, resulting in this larger variance of the validation dataset.

AutoCombat performed non-significantly better on average compared with the other mean and scale adjustment methods (NeuroComBat, CovBat, NeuroHarmonize). However, AutoComBat did nominally reduce the variation of individual features and improve model fit the most, which could be attributed to the strengthened associations between the ASL features and age. Conversely, RELIEF showed the worst performance, which could be attributed to its weakened associations between age and ASL features. These results are similar to those of ASL feature harmonisation, further demonstrating that large ROIs, such as the whole GM or vascular territories, are not strongly affected by the interaction of ASL parameters, age, and measured perfusion. However, these interactions could be more prominently altered in cohorts where CBF and sCoV are affected by pathology (Grade et al., 2015; Gyanwali et al., 2021).

Model performance using all features (T1w, FLAIR, and ASL) did not improve after harmonizing ASL features for most methods. The exception was RELIEF, which showed worse performance in both the validation and testing datasets. This can be attributed to the lower importance of ASL compared to T1w features for estimating brain age, likely due to the high variability of CBF within and between individuals(Clement et al., 2018). However, as a potential earlier biomarker than the structural change detected on T1w-MRI(Grade et al., 2015), ASL-derived CBF as a feature in estimating cerebrovascular ageing might play a greater role in the context of pathology than in healthy ageing.

Harmonisation in the ASL-only model did not change brain age interpretability (i.e., direction of the BAG) for the validation dataset. However, the interpretability changed for the testing datasets with all harmonisation methods resulting in the average BAG being negative, or the brain appearing younger than its chronological age, with RELIEF showing the largest change. This result suggests that harmonisation provides a more truthful image of ageing that is otherwise obscured due to ASL sequence differences, as the Insight 46 cohort on average appeared to be more healthy than its contemporaries(M. B. Dijsselhof et al., 2024; James et al., 2018), and has been estimated to be younger than its chronological age in another brain age study(Wagen et al., 2022). This is, however, difficult to confirm as no brain age studies have been performed in the other datasets used here.

BAG direction did not change in the model using T1w, FLAIR, and ASL after harmonisation, again possibly due to the importance of ASL features in estimating brain age. Expectedly, harmonisation does not remove the inherent regression-to-the-mean effect that creates an age bias in brain age estimation models(de Lange & Cole, 2020). Uncorrected brain age estimations did change brain age interpretability, resulting in large underestimation and worse model accuracy, which can be explained by the older average age of the testing compared to the training datasets. The use of age-bias corrections is debated(de Lange & Cole, 2020), and differences in this study highlight the importance of assessing model performance with and without correction.

### 4.4 Post-hoc analyses

Brain age accuracy in the testing data improved significantly compared with the unharmonised estimations when NeuroCombat was used to harmonise T1w, FLAIR and ASL features. This additionally highlights the higher importance of structural features over ASL features in brain age estimations, possibly due to the large inter-subject variability in CBF(Clement et al., 2018). Compared with the harmonisation of only ASL features and the results of the ASL-only brain age estimations, harmonising all features did have less effect on T1w+FLAIR+ASL brain age estimations. This could be explained by smaller differences between T1w sequences in volumetric brain measurements(van Nederpelt et al., 2023) compared with differences between ASL sequences(Almeida et al., 2018), as measured by intraclass correlation coefficients. The effect of T1w normalisation on the brain age estimation thus clearly highlights the role of feature importance on brain age estimations.

NeuroComBat harmonisation of all features resulted in a positive shift of average BAG (brain appearing older than chronological age). The largest BAG change occurred in the Insight 46 dataset, of which a previous study using T1w volumetric features showed an opposite result with a younger appearing age on average(Wagen et al., 2022). In these cases, where harmonisation changes the BAG interpretation, it is imperative to consider other measures of health and the unharmonised BAG.

The current feature-based approach renders the brain age estimation less sensitive to image-related sequence differences such as susceptibility artefacts or the presence of vascular ASL signal(Alsop et al., 2015). In contrast, deep-learning-based brain age estimations might be more susceptible to these issues, and other methods of image level harmonisations using deep- or transfer-learning approaches(Da-Ano et al., 2021; Hu et al., 2023) might need to be applied before these advanced estimation methods can be used. Additionally, ASL-derived CBF could be altered severely with pathology(Grade et al., 2015), affecting feature distributions. Lastly, many variations in ASL sequences exist that differ in labelling technique, PLD timings, and readout approaches(van Osch et al., 2018), but only a few have been included in this study. Additionally, there is further variety in image processing pipelines(Paschoal et al., 2024), adding to the between-center variability. Therefore, future studies are encouraged to investigate the effect of harmonisation on association of CBF and sCoV in diseased populations, and validate the positive effect of image-level harmonisations in the presence of pathology across a wider range of ASL sequences and processing pipelines.

### 4.5 Limitations

This study has several limitations. First, this study has investigated only a selection of feature-level harmonisation methods, and omitted deep-learning-based harmonisations(Hu et al., 2023) as the former is relatively straightforward to understand and interpret the effect of harmonisation on the feature-based brain age estimation. Although deep-learning methods might harmonise better than feature-level methods, specifically as they might better address spatial variations between ASL sequences, it is more difficult to understand their effect on the harmonisation of ROI-specific CBF and sCoV features. Within these feature-level methods, despite not all publicly available harmonisation methods being tested, the selection covers all groups of methods described by Hu et al.(Hu et al., 2023), assuming that performance will be similar across the group.

The population differences among the testing datasets are another limitation, which make it difficult to disentangle ASL sequence batch-effects from population differences. For example, Insight 46 has a very narrow age range, which complicates harmonisation of datasets with larger age ranges. Although age-matched subsets might offer more insight into the usefulness and correctness of harmonisations, large and homogeneous ASL datasets are not easily available making the studied scenario a realistic example.

Furthermore, many harmonisation methods assume normally distributed features. To avoid non-normality issues in this study, several features (WMH and sCoV) have been log-transformed, however, this decreases the interpretability of the individual measures. In general, and in this study specifically, training datasets suffer from fewer outliers due to the inclusion of healthy participants. However, harmonisation might be less effective in non-normally distributed features and will adversely affect the distribution in the training datasets if testing datasets include patients.

Furthermore, this could result in worse model performance, as seen in the validation results in this study, and this effect could be even more pronounced when harmonising patient data. Therefore, other harmonisation methods that are able to handle non-normal distributions should be studied, especially in the context of pathology. Lastly, the preservation of biological and pathological associations by the harmonisation methods was assessed by determining associations between the ASL features and age in this study. This allows for only a partial view into assessing the (in)ability of the harmonisation methods to disentangle batch and population effects, as shown by the difference of RELIEF compared with the other harmonisation methods in the associations between ASL features and age. Exploring the associations of CBF features and brain age estimations with other health parameters, such as blood pressure and cognitive scores, and perfusion modifiers will allow for a better understanding of whether datasets-specific associations are retained while improving compatibility across datasets.

## 5. Conclusions

In the largest ASL brain age study to date, consisting of several datasets acquired with different ASL sequences and population characteristics, we showed that ASL-derived CBF and sCoV can be harmonised using traditional feature-level harmonisation methods. AutoComBat achieved the highest comparability and model accuracy when considering that latent-factor approaches might remove biological associations. While adding T1w and FLAIR features lowered the effect of harmonisation on brain age estimation performance. The improvement in ASL-feature-only model clearly showed the added value of ASL data harmonisation in multi-cohort ASL analyses, allowing advanced models such as brain age estimations to explore the associations between ageing, cardiovascular risk factors, brain health, and cognitive decline.

## Data and Code Availability

No new data were acquired for this study, and all data has been requested through the respective data request procedures of each study as detailed in the referenced publications. ExploreASL code is freely available at https://github.com/ExploreASL/ExploreASL. Harmonisation and brain age estimation code is under development and is available on request at https://github.com/ExploreASL/cvasl and https://github.com/MDijsselhof/CerebrovascularBrainAge and will be released under an open-source license in the future.

