## Supplemental Files for "Multi-cohort, multi-sequence harmonisation for cerebrovascular brain age"

**Supplementary**

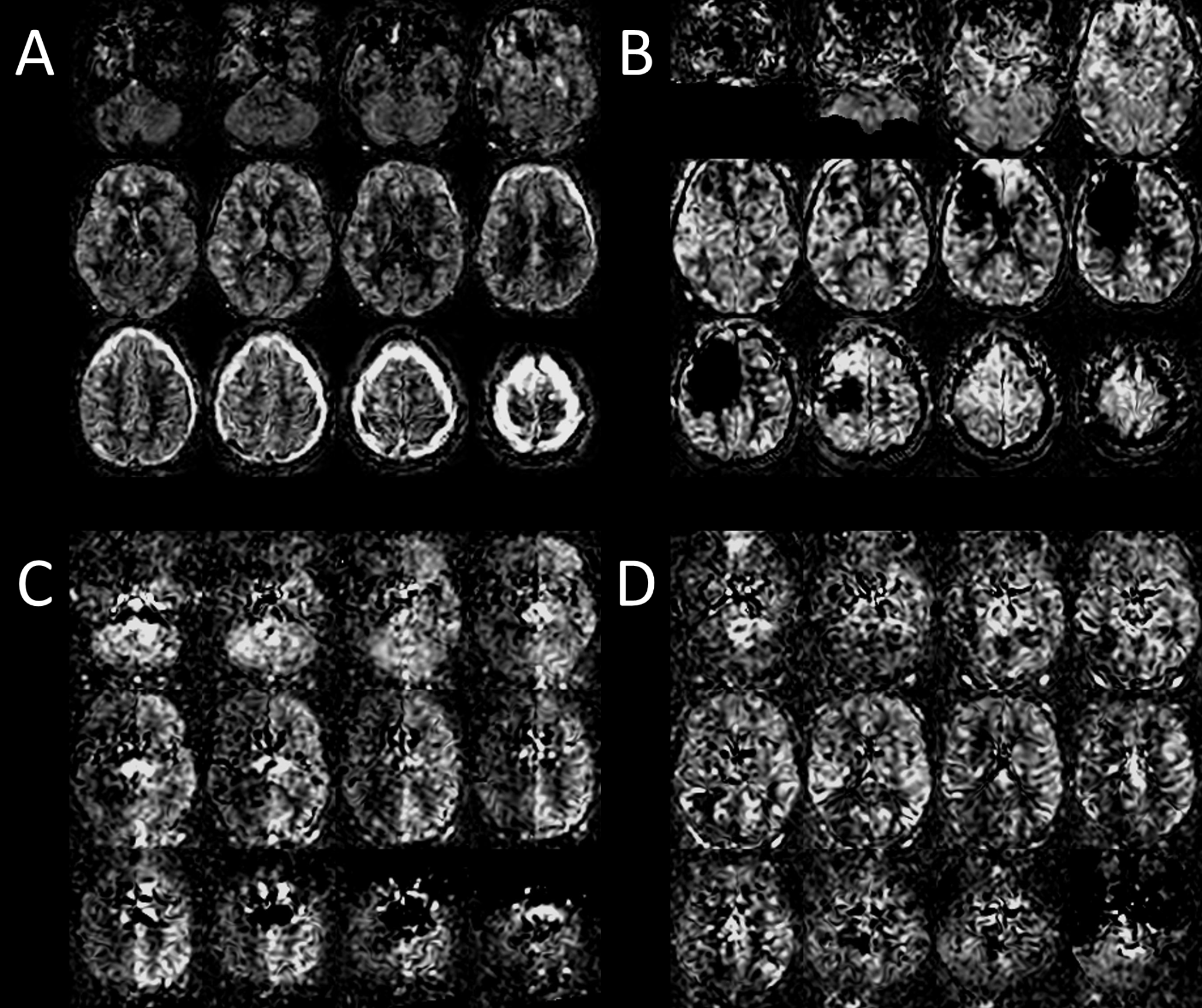

**Supplementary Figure 1:** Examples of excluded CBF maps, showing motion artefacts (A), possible coil issues (B), asymetric labelling efficiency (C), hyperintense vascular signal (C,D), and arterial transit artefact-related hypointense signal (D). *CBF: cerebral blood flow.*

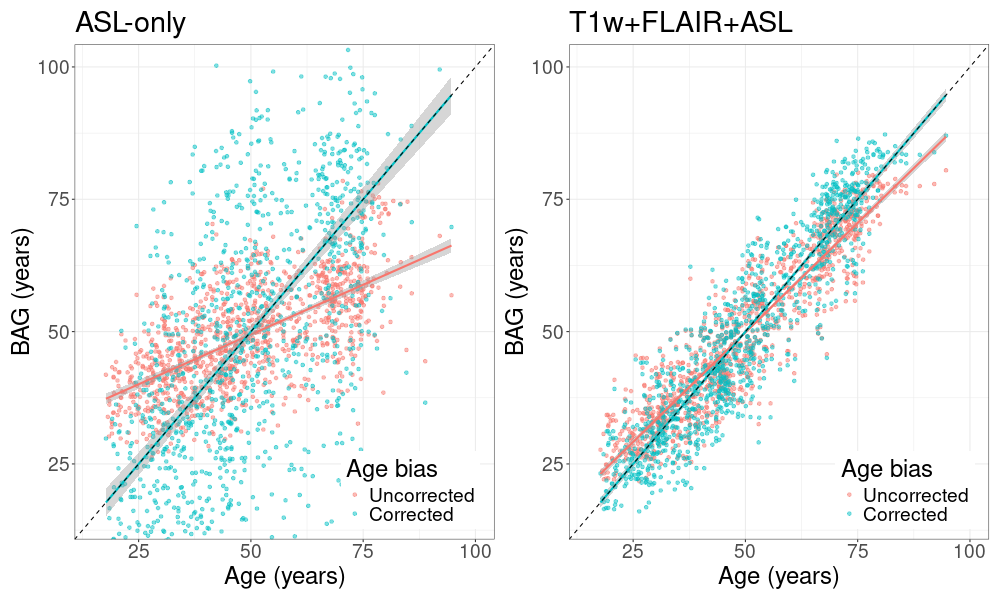

**Supplementary Figure 2:** The association of the validation dataset uncorrected (red) and age bias-corrected (blue) BAGs with the chronological age of the ASL-only and T1w+FLAIR+ASL models before harmonisation. The dashed black line represents a perfect estimation. *ASL: arterial spin labelling; BAG: Brain age gap; FLAIR: Fluid attenuated inversion recovery; T1w: T1-weighted;*

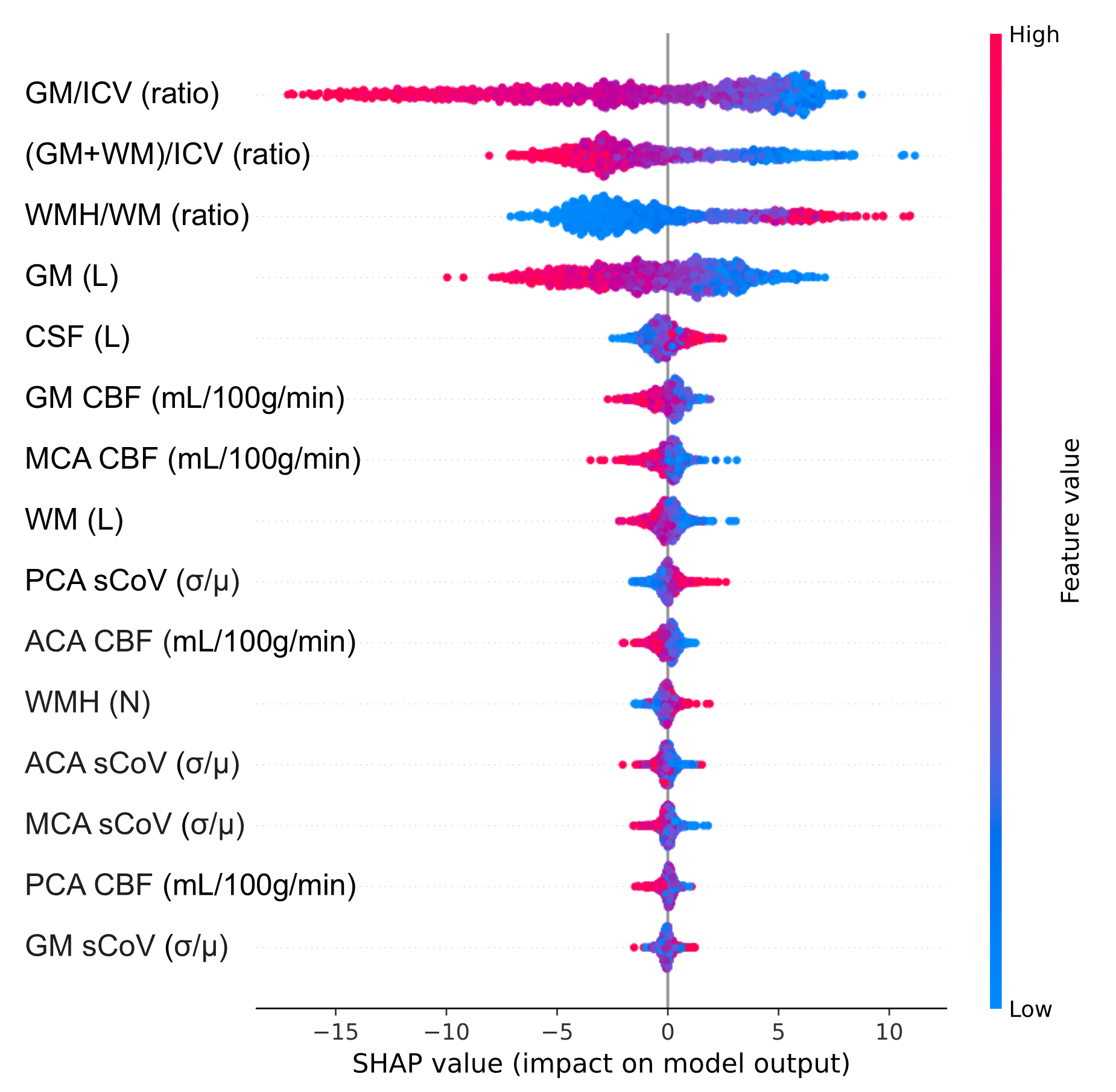

**Supplementary Figure 3:** Feature importance of the T1w+FLAIR+ASL brain age estimation model derived using Shapley values. *ACA: anterior cerebral artery; CBF: cerebral blood flow; CSF: cerebrospinal fluid; GM: grey matter; ICV: intracranial volume; MCA: middle cerebral artery; PCA: posterior cerebral artery; sCoV: spatial coefficient of variation; WM: white matter; WMH: white matter hyperintensities.*

**Supplementary Table 1:** ANCOVA with Tukey post hoc test results, corrected for age and sex, of differences in unharmonised imaging feature between cohort-pairs, shown in mean [confidence interval]. WMH/WM, WMH count, and all sCoV features have been log-transformed. *ACA: anterior cerebral artery; CBF: cerebral blood flow; CSF: cerebrospinal fluid; GM: grey matter; ICV: intracranial volume; MCA: middle cerebral artery; PCA: posterior cerebral artery; sCoV: spatial coefficient of variation; WM: white matter; WMH: white matter hyperintensities.*

| **Feature** | **EDIS - HELIUS** | **EDIS - Insight46** | **EDIS - SABRE** | **EDIS - Training** | **HELIUS - Insight46** | **HELIUS - SABRE** | **HELIUS - Training** | **Insight46 - SABRE** | **Insight46 - Training** | **SABRE - Training** |
| --- | --- | --- | --- | --- | --- | --- | --- | --- | --- | --- |
| **GM (L)** | |  |  |  |  |  |  |  |  |  |
| Difference | -0.04  [-0.05, -0.03] | -0.06  [-0.07, -0.06] | 0  [-0.01, 0.00] | -0.11  [-0.12, -0.10] | -0.02  [-0.03, -0.02] | 0.04  [0.03, 0.04] | -0.07  [-0.08, -0.06] | 0.06  [0.05, 0.07] | -0.05  [-0.05, -0.04] | -0.11  [-0.11, -0.10] |
| P-value | <0.001 | <0.001 | 0.939 | <0.001 | <0.001 | <0.001 | <0.001 | <0.001 | <0.001 | <0.001 |
| **WM (L)** | |  |  |  |  |  |  |  |  |  |
| Difference | -0.08  [-0.09, -0.07] | -0.06  [-0.07, -0.05] | -0.02  [-0.03, -0.01] | -0.10  [-0.11, -0.1] | 0.02  [0.01, 0.03] | 0.06  [0.05, 0.07] | -0.02  [-0.03, -0.02] | 0.04  [0.03, 0.04] | -0.04  [-0.05, -0.04] | -0.08  [-0.09, -0.07] |
| P-value | <0.001 | <0.001 | <0.001 | <0.001 | <0.001 | <0.001 | <0.001 | <0.001 | <0.001 | <0.001 |
| **CSF (L)** | |  |  |  |  |  |  |  |  |  |
| Difference | -0.07  [-0.08, -0.06] | -0.01  [-0.02, 0.00] | 0.01  [0.00, 0.02] | -0.10  [-0.11, -0.09] | 0.06  [0.05, 0.07] | 0.08  [0.07, 0.09] | -0.04  [-0.04, -0.03] | 0.02  [0.01, 0.03] | -0.10  [-0.11, -0.09] | -0.12  [-0.13, -0.11] |
| P-value | <0.001 | 0.785 | 0.021 | <0.001 | <0.001 | <0.001 | <0.001 | <0.001 | <0.001 | <0.001 |
| **GM/ICV (ratio)** | |  |  |  |  |  |  |  |  |  |
| Difference | 0.02  [0.02, 0.03] | -0.01  [-0.01, -0.01] | 0.00  [0.00, 0.00] | 0.01  [0.01, 0.02] | -0.03  [-0.04, -0.03] | -0.02  [-0.03, -0.02] | -0.01  [-0.01, -0.01] | 0.01  [0.01, 0.01] | 0.02  [0.02, 0.03] | 0.01  [0.01, 0.02] |
| P-value | <0.001 | <0.001 | 0.956 | <0.001 | <0.001 | <0.001 | <0.001 | <0.001 | <0.001 | <0.001 |
| **(GM+WM)/ICV ratio** | |  |  |  |  |  |  |  |  |  |
| Difference | 0.01  [0.01, 0.02] | -0.02  [-0.03, -0.01] | -0.01  [-0.02, -0.01] | 0.02  [0.01, 0.02] | -0.03  [-0.04, -0.03] | -0.02  [-0.03, -0.02] | 0.00  [0.00, 0.01] | 0.01  [0.00, 0.01] | 0.04  [0.03, 0.04] | 0.03  [0.02, 0.03] |
| P-value | <0.001 | <0.001 | <0.001 | <0.001 | <0.001 | <0.001 | 0.230 | 0.033 | <0.001 | <0.001 |
| **WMH/WM (ratio)** | |  |  |  |  |  |  |  |  |  |
| Difference | 0.00  [0.00, 0.00] | 0.01  [0.01, 0.01] | 0.00  [0.00, 0.01] | -0.01  [-0.01, -0.01] | 0.01  [0.01, 0.02] | 0.00  [0.00, 0.01] | -0.01  [-0.01, -0.01] | -0.01  [-0.01, -0.01] | -0.02  [-0.03, -0.02] | -0.02  [-0.02, -0.01] |
| P-value | 0.688 | <0.001 | 0.020 | <0.001 | <0.001 | <0.001 | <0.001 | <0.001 | <0.001 | <0.001 |
| **WMH (N)** | |  |  |  |  |  |  |  |  |  |
| Difference | -0.28  [-0.34, -0.21] | 0.05  [-0.02, 0.12] | -0.03  [-0.09, 0.03] | -0.40  [-0.46, -0.33] | 0.33  [0.26, 0.40] | 0.25  [0.19, 0.30] | -0.12  [-0.17, -0.08] | -0.08  [-0.14, -0.02] | -0.45  [-0.52, -0.38] | -0.37  [-0.43, -0.31] |
| P-value | <0.001 | 0.554 | 0.866 | <0.001 | <0.001 | <0.001 | <0.001 | 0.068 | <0.001 | <0.001 |
| **GM CBF (mL/100g/min)** | |  |  |  |  |  |  |  |  |  |
| Difference | 15.84  [14.16, 17.53] | 10.69  [8.88, 12.5] | 23.94  [22.44, 25.45] | 13.89  [12.19, 15.58] | -5.15  [-6.95, -3.36] | 8.10  [6.60, 9.59] | -1.96  [-3.20, -0.71] | 13.25  [11.64, 14.86] | 3.20  [1.38, 5.01] | -10.05  [-11.58, -8.53] |
| P-value | <0.001 | <0.001 | <0.001 | <0.001 | <0.001 | <0.001 | 0.018 | <0.001 | 0.005 | <0.001 |
| **ACA CBF (mL/100g/min)** | |  |  |  |  |  |  |  |  |  |
| Difference | 10.52  [8.45, 12.59] | 5.27  [3.05, 7.50] | 23.57  [21.72, 25.42] | 7.96  [5.87, 10.04] | -5.25  [-7.46, -3.04] | 13.05  [11.21, 14.89] | -2.56  [-4.09, -1.03] | 18.29  [16.32, 20.27] | 2.69  [0.45, 4.92] | -15.61  [-17.48, -13.73] |
| P-value | <0.001 | <0.001 | <0.001 | <0.001 | <0.001 | <0.001 | <0.001 | <0.001 | 0.128 | <0.001 |
| **MCA CBF (mL/100g/min)** | |  |  |  |  |  |  |  |  |  |
| Difference | 18.35  [16.46, 20.24] | 9.80  [7.77, 11.83] | 30.71  [29.02, 32.39] | 15.71  [13.80, 17.61] | -8.55  [-10.56, -6.53] | 12.36  [10.68, 14.04] | -2.64  [-4.04, -1.25] | 20.90  [19.1, 22.71] | 5.90  [3.86, 7.94] | -15.00  [-16.71, -13.29] |
| P-value | <0.001 | <0.001 | <0.001 | <0.001 | <0.001 | <0.001 | 0.002 | <0.001 | <0.001 | <0.001 |
| **PCA CBF (mL/100g/min)** | |  |  |  |  |  |  |  |  |  |
| Difference | 12.69  [10.97, 14.42] | 5.50  [3.65, 7.36] | 18.54  [16.99, 20.08] | 11.08  [9.34, 12.82] | -7.19  [-9.03, -5.35] | 5.84  [4.30, 7.38] | -1.62  [-2.89, -0.34] | 13.03  [11.38, 14.68] | 5.58  [3.71, 7.44] | -7.46  [-9.02, -5.89] |
| P-value | <0.001 | <0.001 | <0.001 | <0.001 | <0.001 | <0.001 | 0.095 | <0.001 | <0.001 | <0.001 |
| **GM sCoV (ratio)** | |  |  |  |  |  |  |  |  |  |
| Difference | 0.40  [0.37, 0.43] | 0.37  [0.34, 0.39] | 0.26  [0.23, 0.28] | 0.92  [0.89, 0.95] | -0.03  [-0.06, -0.01] | -0.15  [-0.17, -0.12] | 0.52  [0.50, 0.54] | -0.11  [-0.14, -0.09] | 0.55  [0.53, 0.58] | 0.67  [0.64, 0.69] |
| P-value | <0.001 | <0.001 | <0.001 | <0.001 | 0.126 | <0.001 | <0.001 | <0.001 | <0.001 | <0.001 |
| **ACA sCoV (ratio)** | |  |  |  |  |  |  |  |  |  |
| Difference | 0.40  [0.38, 0.43] | 0.28  [0.25, 0.31] | 0.21  [0.19, 0.24] | 0.99  [0.96, 1.02] | -0.12  [-0.15, -0.09] | -0.19  [-0.22, -0.17] | 0.58  [0.56, 0.6] | -0.07  [-0.10, -0.05] | 0.70  [0.67, 0.73] | 0.78  [0.75, 0.8] |
| P-value | <0.001 | <0.001 | <0.001 | <0.001 | <0.001 | <0.001 | <0.001 | <0.001 | <0.001 | <0.001 |
| **MCA sCoV (ratio)** | |  |  |  |  |  |  |  |  |  |
| Difference | 0.41  [0.38, 0.44] | 0.49  [0.46, 0.51] | 0.18  [0.16, 0.21] | 0.97  [0.94, 0.99] | 0.07  [0.05, 0.10] | -0.23  [-0.25, -0.21] | 0.56  [0.54, 0.58] | -0.3  [-0.33, -0.28] | 0.48  [0.45, 0.51] | 0.79  [0.76, 0.81] |
| P-value | <0.001 | <0.001 | <0.001 | <0.001 | <0.001 | <0.001 | <0.001 | <0.001 | <0.001 | <0.001 |
| **PCA sCoV (ratio)** | |  |  |  |  |  |  |  |  |  |
| Difference | 0.51  [0.47, 0.54] | 0.49  [0.46, 0.52] | 0.42  [0.39, 0.44] | 1.09  [1.06, 1.12] | -0.02  [-0.05, 0.02] | -0.09  [-0.12, -0.06] | 0.59  [0.56, 0.61] | -0.07  [-0.1, -0.04] | 0.60  [0.57, 0.64] | 0.68  [0.65, 0.71] |
| P-value | <0.001 | <0.001 | <0.001 | <0.001 | <0.001 | <0.001 | <0.001 | <0.001 | <0.001 | <0.001 |

**Supplementary Table 2**: GM, ACA, MCA, and PCA CBF and log-transformed sCoV of the combined or separate testing cohort-pairs per harmonisation method. *ACA: anterior cerebral artery; CBF: cerebral blood flow; GM: grey matter; MCA: middle cerebral artery; PCA: posterior cerebral artery; sCoV: spatial coefficient of variation.*

| **Harmonisation method** | | **GM CBF (mL/100g/min)** | **ACA CBF (mL/100g/min)** | **MCA CBF (mL/100g/min)** | **PCA CBF (mL/100g/min)** | **GM sCoV (ratio)** | **ACA sCoV (ratio)** | **MCA sCoV (ratio)** | **PCA sCoV (ratio)** |
| --- | --- | --- | --- | --- | --- | --- | --- | --- | --- |
| Unharmonised | | 61.48 ± 14.46 | 75.88 ± 17.25 | 68.29 ± 16.87 | 54.69 ± 13.73 | -1.04 ± 0.41 | -1.17 ± 0.45 | -1.08 ± 0.42 | -1.08 ± 0.46 |
| NeuroComBat | | 61.48 ± 12.66 | 75.88 ± 15.01 | 68.29 ± 13.97 | 54.69 ± 12.64 | -1.04 ± 0.19 | -1.17 ± 0.20 | -1.08 ± 0.19 | -1.08 ± 0.22 |
| CovBat | | 61.48 ± 12.66 | 75.88 ± 15.01 | 68.29 ± 13.97 | 54.69 ± 12.65 | -1.04 ± 0.19 | -1.17 ± 0.20 | -1.08 ± 0.19 | -1.08 ± 0.22 |
| NeuroHarmonize | | 61.45 ± 12.48 | 75.85 ± 14.93 | 68.27 ± 13.90 | 54.67 ± 12.43 | -1.04 ± 0.19 | -1.17 ± 0.20 | -1.08 ± 0.19 | -1.07 ± 0.22 |
| OPN ComBat | | 61.40 ± 12.45 | 75.77 ± 14.87 | 68.20 ± 13.86 | 54.63 ± 12.41 | -1.04 ± 0.19 | -1.17 ± 0.20 | -1.08 ± 0.19 | -1.08 ± 0.22 |
| AutoComBat | | 61.92 ± 12.83 | 76.19 ± 15.14 | 68.78 ± 14.24 | 55.1 ± 12.70 | -1.04 ± 0.19 | -1.17 ± 0.20 | -1.08 ± 0.19 | -1.09 ± 0.22 |
| RELIEF | | 61.47 ± 10.98 | 75.86 ± 13.32 | 68.27 ± 12.16 | 54.68 ± 10.91 | -1.04 ± 0.12 | -1.17 ± 0.13 | -1.08 ± 0.10 | -1.08 ± 0.15 |
| **Cohort** | **Harmonisation method** | **GM CBF (mL/100g/min)** | **ACA CBF (mL/100g/min)** | **MCA CBF (mL/100g/min)** | **PCA CBF (mL/100g/min)** | **GM sCoV (ratio)** | **ACA sCoV (ratio)** | **MCA sCoV (ratio)** | **PCA sCoV (ratio)** |
| Training | Unharmonised | 66.2 ± 11.29 | 82.41 ± 13.34 | 73.74 ± 12.07 | 58.15 ± 10.64 | -1.48 ± 0.11 | -1.67 ± 0.12 | -1.53 ± 0.08 | -1.57 ± 0.13 |
|  | NeuroComBat | 65.65 ± 12.98 | 79.76 ± 15.31 | 72.45 ± 14.26 | 58.17 ± 12.8 | -1.07 ± 0.18 | -1.2 ± 0.2 | -1.09 ± 0.18 | -1.12 ± 0.21 |
|  | CovBat | 65.65 ± 12.98 | 79.76 ± 15.3 | 72.45 ± 14.26 | 58.17 ± 12.8 | -1.07 ± 0.18 | -1.2 ± 0.2 | -1.09 ± 0.18 | -1.12 ± 0.21 |
|  | NeuroHarmonize | 65.33 ± 12.9 | 79.81 ± 15.26 | 72.19 ± 14.25 | 57.49 ± 12.71 | -1.05 ± 0.19 | -1.18 ± 0.2 | -1.07 ± 0.18 | -1.09 ± 0.21 |
|  | OPN ComBat | 64.93 ± 12.83 | 79.29 ± 15.11 | 72.05 ± 14.07 | 57.07 ± 12.61 | -1.06 ± 0.17 | -1.19 ± 0.18 | -1.09 ± 0.17 | -1.12 ± 0.2 |
|  | AutoComBat | 66.18 ± 13.45 | 80.23 ± 15.63 | 73.26 ± 14.8 | 58.33 ± 13.19 | -1.08 ± 0.19 | -1.19 ± 0.2 | -1.09 ± 0.19 | -1.14 ± 0.22 |
|  | RELIEF | 61.48 ± 10.45 | 75.88 ± 12.42 | 68.29 ± 11.4 | 54.69 ± 9.86 | -1.04 ± 0.11 | -1.17 ± 0.11 | -1.08 ± 0.08 | -1.08 ± 0.13 |
| HELIUS | Unharmonised | 61.27 ± 9.88 | 77.07 ± 12.73 | 68.03 ± 11.3 | 54.27 ± 9.84 | -0.94 ± 0.14 | -1.07 ± 0.13 | -0.97 ± 0.12 | -0.95 ± 0.19 |
|  | NeuroComBat | 62.71 ± 11.6 | 77.02 ± 14.17 | 69.42 ± 12.95 | 55.94 ± 12.06 | -1.05 ± 0.19 | -1.18 ± 0.2 | -1.08 ± 0.19 | -1.09 ± 0.23 |
|  | CovBat | 62.71 ± 11.58 | 77.02 ± 14.13 | 69.42 ± 12.92 | 55.94 ± 12.05 | -1.05 ± 0.19 | -1.18 ± 0.2 | -1.08 ± 0.19 | -1.09 ± 0.23 |
|  | NeuroHarmonize | 61.56 ± 11.53 | 75.87 ± 14.16 | 69.31 ± 12.95 | 55.22 ± 12.01 | -1.06 ± 0.19 | -1.19 ± 0.19 | -1.09 ± 0.18 | -1.09 ± 0.22 |
|  | OPN ComBat | 61.61 ± 11.53 | 75.95 ± 14.05 | 68.46 ± 12.83 | 54.6 ± 12.02 | -1.04 ± 0.19 | -1.17 ± 0.19 | -1.07 ± 0.18 | -1.08 ± 0.23 |
|  | AutoComBat | 62.59 ± 11.7 | 76.93 ± 14.22 | 69.4 ± 13.03 | 55.48 ± 12.2 | -1.04 ± 0.2 | -1.17 ± 0.2 | -1.07 ± 0.19 | -1.09 ± 0.23 |
|  | RELIEF | 61.48 ± 10.06 | 75.88 ± 12.85 | 68.29 ± 11.28 | 54.69 ± 10.37 | -1.04 ± 0.11 | -1.17 ± 0.12 | -1.08 ± 0.1 | -1.08 ± 0.15 |
| SABRE | Unharmonised | 47.76 ± 11.47 | 58.98 ± 13.1 | 50.38 ± 12.32 | 43.63 ± 12.96 | -0.74 ± 0.24 | -0.83 ± 0.25 | -0.71 ± 0.25 | -0.8 ± 0.29 |
|  | NeuroComBat | 57.26 ± 11.6 | 71.95 ± 14.22 | 64.1 ± 13.09 | 51.12 ± 11.89 | -1 ± 0.19 | -1.14 ± 0.2 | -1.06 ± 0.19 | -1.03 ± 0.22 |
|  | CovBat | 57.26 ± 11.62 | 71.95 ± 14.25 | 64.1 ± 13.12 | 51.12 ± 11.9 | -1 ± 0.19 | -1.14 ± 0.2 | -1.06 ± 0.19 | -1.03 ± 0.22 |
|  | NeuroHarmonize | 57.94 ± 11.52 | 72.3 ± 14.1 | 64.39 ± 12.98 | 52.21 ± 11.78 | -1.02 ± 0.19 | -1.15 ± 0.2 | -1.07 ± 0.19 | -1.06 ± 0.22 |
|  | OPN ComBat | 57.92 ± 11.32 | 72.38 ± 13.88 | 64.44 ± 12.84 | 52.14 ± 11.67 | -1.01 ± 0.19 | -1.15 ± 0.2 | -1.06 ± 0.19 | -1.04 ± 0.22 |
|  | AutoComBat | 57.2 ± 11.56 | 71.52 ± 14.15 | 63.8 ± 13.04 | 51.64 ± 11.87 | -1.02 ± 0.19 | -1.16 ± 0.2 | -1.07 ± 0.19 | -1.05 ± 0.22 |
|  | RELIEF | 61.48 ± 8.93 | 75.88 ± 10.97 | 68.29 ± 9.9 | 54.69 ± 8.89 | -1.04 ± 0.09 | -1.17 ± 0.1 | -1.08 ± 0.08 | -1.08 ± 0.12 |
| EDIS | Unharmonised | 72.33 ± 15.25 | 83.15 ± 18.56 | 81.78 ± 17.39 | 62.59 ± 14.88 | -0.49 ± 0.27 | -0.63 ± 0.29 | -0.53 ± 0.28 | -0.4 ± 0.29 |
|  | NeuroComBat | 57.9 ± 11.33 | 72.55 ± 13.97 | 64.79 ± 12.79 | 51.54 ± 11.64 | -1 ± 0.18 | -1.14 ± 0.2 | -1.06 ± 0.19 | -1.04 ± 0.22 |
|  | CovBat | 57.9 ± 11.35 | 72.55 ± 14 | 64.79 ± 12.8 | 51.54 ± 11.68 | -1 ± 0.18 | -1.14 ± 0.2 | -1.06 ± 0.19 | -1.04 ± 0.22 |
|  | NeuroHarmonize | 57.94 ± 11.3 | 72.26 ± 13.91 | 65.07 ± 12.68 | 52.38 ± 11.52 | -1.03 ± 0.18 | -1.16 ± 0.2 | -1.08 ± 0.19 | -1.07 ± 0.21 |
|  | OPN ComBat | 59.56 ± 11.24 | 74.05 ± 13.82 | 66.11 ± 12.64 | 53.69 ± 11.51 | -1.03 ± 0.19 | -1.17 ± 0.2 | -1.08 ± 0.19 | -1.06 ± 0.21 |
|  | AutoComBat | 59.18 ± 11.25 | 73.64 ± 13.83 | 65.86 ± 12.71 | 52.88 ± 11.43 | -1.02 ± 0.19 | -1.16 ± 0.2 | -1.07 ± 0.19 | -1.05 ± 0.21 |
|  | RELIEF | 61.48 ± 14.54 | 75.88 ± 17.32 | 68.29 ± 16.19 | 54.69 ± 14.64 | -1.04 ± 0.17 | -1.17 ± 0.18 | -1.08 ± 0.15 | -1.08 ± 0.21 |
| Insight46 | Unharmonised | 61.16 ± 14.79 | 77.42 ± 18.7 | 71.45 ± 17.84 | 56.76 ± 15.36 | -0.86 ± 0.22 | -0.91 ± 0.25 | -1.01 ± 0.23 | -0.88 ± 0.24 |
|  | NeuroComBat | 57.16 ± 11.44 | 71.88 ± 14.14 | 64.2 ± 13.04 | 50.78 ± 11.67 | -1 ± 0.18 | -1.15 ± 0.19 | -1.06 ± 0.18 | -1.03 ± 0.21 |
|  | CovBat | 57.16 ± 11.44 | 71.89 ± 14.12 | 64.2 ± 13.01 | 50.79 ± 11.66 | -1 ± 0.18 | -1.15 ± 0.19 | -1.06 ± 0.18 | -1.03 ± 0.21 |
|  | NeuroHarmonize | 58.57 ± 11.56 | 73.04 ± 14.24 | 64.25 ± 13.33 | 51.05 ± 11.93 | -1.01 ± 0.18 | -1.16 ± 0.19 | -1.08 ± 0.18 | -1.04 ± 0.21 |
|  | OPN ComBat | 58.16 ± 12.12 | 72.71 ± 15.02 | 64.96 ± 13.98 | 52.24 ± 12.42 | -1.02 ± 0.19 | -1.16 ± 0.2 | -1.07 ± 0.19 | -1.04 ± 0.21 |
|  | AutoComBat | 58.29 ± 11.42 | 72.97 ± 14.21 | 65.37 ± 13.23 | 52.24 ± 11.72 | -1.01 ± 0.18 | -1.16 ± 0.19 | -1.07 ± 0.18 | -1.04 ± 0.2 |
|  | RELIEF | 61.14 ± 13.21 | 75.53 ± 16.1 | 68.08 ± 14.95 | 54.21 ± 13.69 | -1.04 ± 0.15 | -1.17 ± 0.16 | -1.08 ± 0.13 | -1.07 ± 0.19 |

**Supplementary Table 3**: Regression results for the associations of age with CBF and log-transformed sCoV. *ACA: anterior cerebral artery; CBF: cerebral blood flow; CI: confidence interval; GM: grey matter; MCA: middle cerebral artery; PCA: posterior cerebral artery; sCoV: spatial coefficient of variation.*

| **CBF (mL/100g/min)** | **Method** | **Beta** | **CI** | **R^2^** | **P-value** |
| --- | --- | --- | --- | --- | --- |
| GM | Unharmonised | -0.37 | -0.41, -0.34 | 0.13 | < 0.001 |
|  | NeuroComBat | -0.45 | -0.49, -0.41 | 0.15 | < 0.001 |
|  | CovBat | -0.45 | -0.49, -0.41 | 0.15 | < 0.001 |
|  | NeuroHarmonize | -0.43 | -0.47, -0.39 | 0.13 | < 0.001 |
|  | OPNComBat | -0.42 | -0.46, -0.38 | 0.12 | < 0.001 |
|  | AutoComBat | -0.47 | -0.51, -0.44 | 0.17 | < 0.001 |
|  | RELIEF | -0.28 | -0.33, -0.23 | 0.04 | < 0.001 |
| ACA | Unharmonised | -0.31 | -0.34, -0.28 | 0.13 | < 0.001 |
|  | NeuroComBat | -0.30 | -0.34, -0.27 | 0.09 | < 0.001 |
|  | CovBat | -0.30 | -0.34, -0.27 | 0.09 | < 0.001 |
|  | NeuroHarmonize | -0.30 | -0.33, -0.26 | 0.09 | < 0.001 |
|  | OPNComBat | -0.28 | -0.32, -0.25 | 0.08 | < 0.001 |
|  | AutoComBat | -0.32 | -0.36, -0.29 | 0.11 | < 0.001 |
|  | RELIEF | -0.17 | -0.21, -0.13 | 0.02 | < 0.001 |
| MCA | Unharmonised | -0.31 | -0.34, -0.28 | 0.12 | < 0.001 |
|  | NeuroComBat | -0.39 | -0.42, -0.35 | 0.13 | < 0.001 |
|  | CovBat | -0.39 | -0.42, -0.35 | 0.13 | < 0.001 |
|  | NeuroHarmonize | -0.38 | -0.41, -0.34 | 0.13 | < 0.001 |
|  | OPNComBat | -0.37 | -0.41, -0.33 | 0.12 | < 0.001 |
|  | AutoComBat | -0.41 | -0.45, -0.38 | 0.16 | < 0.001 |
|  | RELIEF | -0.24 | -0.29, -0.2 | 0.04 | < 0.001 |
| PCA | Unharmonised | -0.29 | -0.33, -0.25 | 0.07 | < 0.001 |
|  | NeuroComBat | -0.34 | -0.39, -0.3 | 0.09 | < 0.001 |
|  | CovBat | -0.34 | -0.38, -0.3 | 0.09 | < 0.001 |
|  | NeuroHarmonize | -0.31 | -0.35, -0.27 | 0.07 | < 0.001 |
|  | OPNComBat | -0.29 | -0.33, -0.25 | 0.06 | < 0.001 |
|  | AutoComBat | -0.35 | -0.39, -0.31 | 0.09 | < 0.001 |
|  | RELIEF | -0.19 | -0.24, -0.14 | 0.02 | < 0.001 |
| **sCoV (ratio)** | **Method** | **Beta** | **CI** | **R^2^** | **P-value** |
| GM | Unharmonised | 20.53 | 19.44, 21.61 | 0.32 | < 0.001 |
|  | NeuroComBat | 14.00 | 11.16, 16.84 | 0.03 | < 0.001 |
|  | CovBat | 13.97 | 11.14, 16.81 | 0.03 | < 0.001 |
|  | NeuroHarmonize | 10.77 | 7.89, 13.65 | 0.02 | < 0.001 |
|  | OPNComBat | 11.54 | 8.6, 14.48 | 0.02 | < 0.001 |
|  | AutoComBat | 15.96 | 13.17, 18.76 | 0.04 | < 0.001 |
|  | RELIEF | 13.28 | 8.78, 17.77 | 0.01 | < 0.001 |
| ACA | Unharmonised | 18.75 | 17.75, 19.74 | 0.32 | < 0.001 |
|  | NeuroComBat | 8.74 | 6.03, 11.46 | 0.01 | < 0.001 |
|  | CovBat | 8.72 | 6.01, 11.43 | 0.01 | < 0.001 |
|  | NeuroHarmonize | 6.92 | 4.19, 9.65 | 0.01 | < 0.001 |
|  | OPNComBat | 6.62 | 3.81, 9.42 | 0.01 | < 0.001 |
|  | AutoComBat | 8.36 | 5.66, 11.05 | 0.01 | < 0.001 |
|  | RELIEF | 5.41 | 1.19, 9.63 | 0.00 | 0.012 |
| MCA | Unharmonised | 18.91 | 17.84, 19.98 | 0.29 | < 0.001 |
|  | NeuroComBat | 4.43 | 1.52, 7.34 | 0.00 | 0.003 |
|  | CovBat | 4.41 | 1.5, 7.31 | 0.00 | 0.003 |
|  | NeuroHarmonize | 2.08 | -0.85, 5.01 | 0.00 | 0.163 |
|  | OPNComBat | 2.96 | -0.05, 5.98 | 0.00 | 0.054 |
|  | AutoComBat | 4.58 | 1.68, 7.48 | 0.00 | 0.002 |
|  | RELIEF | 5.73 | 0.28, 11.19 | 0.00 | 0.040 |
| PCA | Unharmonised | 18.02 | 17.06, 18.98 | 0.32 | < 0.001 |
|  | NeuroComBat | 14.07 | 11.66, 16.49 | 0.04 | < 0.001 |
|  | CovBat | 14.05 | 11.64, 16.46 | 0.04 | < 0.001 |
|  | NeuroHarmonize | 10.95 | 8.48, 13.43 | 0.03 | < 0.001 |
|  | OPNComBat | 13.87 | 11.39, 16.35 | 0.04 | < 0.001 |
|  | AutoComBat | 18.85 | 16.5, 21.2 | 0.08 | < 0.001 |
|  | RELIEF | 13.33 | 9.74, 16.92 | 0.02 | < 0.001 |

**Supplementary Table 4:** BAG and MAE of the testing cohort-pairs separately per harmonisation method, obtained using ASL-only (A) features or T1w+FLAIR+ASL (B) features to predict brain age. *ASL: arterial spin labelling; BAG: Brain age gap; FLAIR: Fluid attenuated inversion recovery; MAE: mean absolute error; T1w: T1-weighted.*

| A. Cohort | Method | BAG  (μ ± σ) | MAE  (μ ± σ) |
| --- | --- | --- | --- |
| Validation | Unharmonised | -0.23 ± 13.03 | 10.65 ± 7.5 |
|  | NeuroComBat | -0.43 ± 13.88 | 11.32 ± 8.04 |
|  | CovBat | -0.22 ± 13.74 | 11.19 ± 7.96 |
|  | NeuroHarmonize | -0.24 ± 13.90 | 11.37 ± 8.00 |
|  | OPNested ComBat | -0.12 ± 13.62 | 11.15 ± 7.81 |
|  | AutoComBat | -0.18 ± 13.06 | 10.62 ± 7.61 |
|  | RELIEF | -0.22 ± 12.91 | 10.42 ± 7.62 |
| HELIUS | Unharmonised | 1.86 ± 10.89 | 9.24 ± 6.04 |
|  | NeuroComBat | -0.89 ± 8.23 | 6.68 ± 4.88 |
|  | CovBat | -1.04 ± 8.14 | 6.59 ± 4.88 |
|  | NeuroHarmonize | -0.33 ± 8.01 | 6.54 ± 4.63 |
|  | OPNested ComBat | -0.66 ± 8.32 | 6.64 ± 5.05 |
|  | AutoComBat | -0.29 ± 8.35 | 6.67 ± 5.02 |
|  | RELIEF | -2.22 ± 8.82 | 7.4 ± 5.27 |
| SABRE | Unharmonised | 8.91 ± 6.06 | 9.73 ± 4.65 |
|  | NeuroComBat | -2.36 ± 7.23 | 6.07 ± 4.58 |
|  | CovBat | -2.75 ± 7.18 | 6.15 ± 4.62 |
|  | NeuroHarmonize | -2.85 ± 6.98 | 5.99 ± 4.57 |
|  | OPNested ComBat | -2.91 ± 7.31 | 6.16 ± 4.89 |
|  | AutoComBat | -1.46 ± 7.15 | 5.73 ± 4.51 |
|  | RELIEF | -8.98 ± 7.51 | 9.93 ± 6.19 |
| EDIS | Unharmonised | -13.72 ± 13.41 | 16.3 ± 10.10 |
|  | NeuroComBat | -1.85 ± 8.03 | 6.53 ± 5.02 |
|  | CovBat | -2.4 ± 7.96 | 6.65 ± 4.98 |
|  | NeuroHarmonize | -2.53 ± 7.73 | 6.44 ± 4.95 |
|  | OPNested ComBat | -2.82 ± 8.26 | 6.91 ± 5.32 |
|  | AutoComBat | -1.61 ± 8.21 | 6.64 ± 5.07 |
|  | RELIEF | -4.1 ± 10.33 | 8.78 ± 6.79 |
| Insight 46 | Unharmonised | -3.09 ± 13.7 | 11.25 ± 8.38 |
|  | NeuroComBat | -0.4 ± 8.30 | 6.39 ± 5.30 |
|  | CovBat | -0.8 ± 8.30 | 6.48 ± 5.23 |
|  | NeuroHarmonize | -1.04 ± 8.33 | 6.51 ± 5.28 |
|  | OPNested ComBat | -1.07 ± 8.74 | 6.78 ± 5.59 |
|  | AutoComBat | -0.71 ± 8.29 | 6.56 ± 5.11 |
|  | RELIEF | -3.05 ± 10.34 | 8.78 ± 6.24 |
| B. Cohort | Method | BAG  (μ ± σ) | MAE  (μ ± σ) |
| Validation | Unharmonised | -0.07 ± 6.47 | 5.11 ± 3.98 |
|  | NeuroComBat | 0.11 ± 6.69 | 5.33 ± 4.03 |
|  | CovBat | -0.08 ± 6.52 | 5.13 ± 4.01 |
|  | NeuroHarmonize | 0.11 ± 6.78 | 5.39 ± 4.11 |
|  | OPNested ComBat | -0.06 ± 6.56 | 5.15 ± 4.06 |
|  | AutoComBat | -0.05 ± 6.5 | 5.12 ± 4 |
|  | RELIEF | -0.05 ± 6.61 | 5.21 ± 4.06 |
|  | NeuroComBat (all features) | 0.00 ± 7.3 | 5.77 ± 4.47 |
| HELIUS | Unharmonised | 2.56 ± 6.8 | 5.76 ± 4.43 |
|  | NeuroComBat | 1.84 ± 7.2 | 5.83 ± 4.61 |
|  | CovBat | 2.25 ± 7.16 | 5.93 ± 4.59 |
|  | NeuroHarmonize | 1.85 ± 7.19 | 5.89 ± 4.52 |
|  | OPNested ComBat | 2.24 ± 7.12 | 5.9 ± 4.56 |
|  | AutoComBat | 2.23 ± 7.04 | 5.82 ± 4.54 |
|  | RELIEF | 1.86 ± 7.2 | 5.88 ± 4.55 |
|  | NeuroComBat (all features) | 4.04 ± 7.18 | 6.74 ± 4.72 |
| SABRE | Unharmonised | -2.98 ± 6.35 | 5.42 ± 4.44 |
|  | NeuroComBat | -3.53 ± 7.13 | 6.02 ± 5.2 |
|  | CovBat | -3.04 ± 6.99 | 5.72 ± 5.03 |
|  | NeuroHarmonize | -3.46 ± 7.12 | 6.01 ± 5.14 |
|  | OPNested ComBat | -3.29 ± 7.02 | 5.86 ± 5.08 |
|  | AutoComBat | -3.11 ± 6.95 | 5.74 ± 5.01 |
|  | RELIEF | -3.75 ± 7.33 | 6.21 ± 5.4 |
|  | NeuroComBat (all features) | 3.06 ± 4.66 | 4.52 ± 3.27 |
| EDIS | Unharmonised | -1.25 ± 7.19 | 5.62 ± 4.64 |
|  | NeuroComBat | 0.16 ± 7.52 | 5.92 ± 4.63 |
|  | CovBat | 0.6 ± 7.53 | 5.92 ± 4.68 |
|  | NeuroHarmonize | 0.31 ± 7.55 | 5.96 ± 4.63 |
|  | OPNested ComBat | 0.29 ± 7.59 | 5.99 ± 4.66 |
|  | AutoComBat | 0.52 ± 7.55 | 5.95 ± 4.66 |
|  | RELIEF | -0.16 ± 7.79 | 6.15 ± 4.77 |
|  | NeuroComBat (all features) | 3.6 ± 5.21 | 5.21 ± 3.6 |
| Insight46 | Unharmonised | -6.94 ± 5.61 | 7.46 ± 4.89 |
|  | NeuroComBat | -6.82 ± 6.26 | 7.51 ± 5.41 |
|  | CovBat | -6.35 ± 6.23 | 7.1 ± 5.35 |
|  | NeuroHarmonize | -6.84 ± 6.51 | 7.6 ± 5.6 |
|  | OPNested ComBat | -6.67 ± 6.55 | 7.46 ± 5.62 |
|  | AutoComBat | -6.44 ± 6.24 | 7.24 ± 5.29 |
|  | RELIEF | -7.55 ± 6.68 | 8.24 ± 5.82 |
|  | NeuroComBat (all features) | 3.72 ± 3.69 | 4.55 ± 2.6 |

**Supplementary Table 5:** BAG and MAE differences between harmonisation methods in the testing dataset, for ASL-only (A) and T1w+FLAIR+ASL features (B). *ASL: arterial spin labelling; BAG: Brain age gap; CI: confidence interval; FLAIR: Fluid attenuated inversion recovery; MAE: mean absolute error; T1w: T1-weighted.*

| **ASL-only** | **BAG difference (mean [CI])** | **P-value** | **MAE difference (mean [CI])** | **P-value** |
| --- | --- | --- | --- | --- |
| NeuroComBat - Unharmonised | -2.13 [-3.02, -1.23] | < 0.001 | -4.69 [-5.24, -4.15] | < 0.001 |
| CovBat - Unharmonised | -2.48 [-3.37, -1.59] | < 0.001 | -4.66 [-5.2, -4.12] | < 0.001 |
| NeuroHarmonize - Unharmonised | -2.37 [-3.26, -1.48] | < 0.001 | -4.76 [-5.31, -4.22] | < 0.001 |
| OPNested ComBat - Unharmonised | -2.55 [-3.44, -1.65] | < 0.001 | -4.54 [-5.08, -4] | < 0.001 |
| AutoComBat - Unharmonised | -1.63 [-2.52, -0.74] | < 0.001 | -4.77 [-5.31, -4.23] | < 0.001 |
| RELIEF - Unharmonised | -5.72 [-6.62, -4.83] | < 0.001 | -2.3 [-2.84, -1.76] | < 0.001 |
| CovBat - NeuroComBat | -0.35 [-1.24, 0.54] | 0.909 | 0.04 [-0.51, 0.58] | 1.000 |
| NeuroHarmonize - NeuroComBat | -0.24 [-1.13, 0.65] | 0.985 | -0.07 [-0.61, 0.47] | 1.000 |
| OPNested ComBat - NeuroComBat | -0.42 [-1.31, 0.47] | 0.812 | 0.15 [-0.39, 0.7] | 0.981 |
| AutoComBat - NeuroComBat | 0.5 [-0.4, 1.39] | 0.655 | -0.08 [-0.62, 0.47] | 1.000 |
| RELIEF - NeuroComBat | -3.59 [-4.49, -2.7] | < 0.001 | 2.39 [1.85, 2.94] | < 0.001 |
| NeuroHarmonize - CovBat | 0.11 [-0.78, 1] | 1.000 | -0.11 [-0.65, 0.44] | 0.998 |
| OPNested ComBat - CovBat | -0.07 [-0.96, 0.83] | 1.000 | 0.12 [-0.43, 0.66] | 0.995 |
| AutoComBat - CovBat | 0.85 [-0.04, 1.74] | 0.076 | -0.11 [-0.66, 0.43] | 0.997 |
| RELIEF - CovBat | -3.24 [-4.14, -2.35] | < 0.001 | 2.36 [1.82, 2.9] | < 0.001 |
| OPNested ComBat - NeuroHarmonize | -0.18 [-1.07, 0.72] | 0.997 | 0.22 [-0.32, 0.77] | 0.889 |
| AutoComBat - NeuroHarmonize | 0.74 [-0.16, 1.63] | 0.183 | -0.01 [-0.55, 0.54] | 1.000 |
| RELIEF - NeuroHarmonize | -3.35 [-4.25, -2.46] | < 0.001 | 2.46 [1.92, 3.01] | < 0.001 |
| AutoComBat - OPNested ComBat | 0.92 [0.02, 1.81] | 0.040 | -0.23 [-0.77, 0.31] | 0.875 |
| RELIEF - OPNested ComBat | -3.18 [-4.07, -2.28] | < 0.001 | 2.24 [1.7, 2.78] | < 0.001 |
| RELIEF - AutoComBat | -4.09 [-4.98, -3.2] | < 0.001 | 2.47 [1.93, 3.01] | < 0.001 |
| **T1w+FLAIR+ASL** | **BAG difference (mean [CI])** | **P-value** | **MAE difference (mean [CI])** | **P-value** |
| NeuroComBat - Unharmonised | -0.12 [-0.87, 0.63] | 1.000 | 0.3 [-0.19, 0.79] | 0.570 |
| NeuroComBat (all features) - Unharmonised | 5.19 [4.44, 5.94] | < 0.001 | -0.57 [-1.05, -0.08] | 0.009 |
| CovBat - Unharmonised | 0.34 [-0.42, 1.09] | 0.880 | 0.16 [-0.33, 0.65] | 0.975 |
| NeuroHarmonize - Unharmonised | -0.06 [-0.82, 0.69] | 1.000 | 0.34 [-0.15, 0.82] | 0.418 |
| OPNested ComBat - Unharmonised | 0.14 [-0.62, 0.89] | 0.999 | 0.27 [-0.22, 0.75] | 0.711 |
| AutoComBat - Unharmonised | 0.27 [-0.48, 1.03] | 0.957 | 0.16 [-0.33, 0.64] | 0.976 |
| RELIEF - Unharmonised | -0.37 [-1.12, 0.39] | 0.822 | 0.54 [0.06, 1.03] | 0.017 |
| NeuroComBat (all features) - NeuroComBat | 5.31 [4.56, 6.06] | < 0.001 | -0.87 [-1.35, -0.38] | < 0.001 |
| CovBat - NeuroComBat | 0.46 [-0.3, 1.21] | 0.599 | -0.14 [-0.63, 0.35] | 0.988 |
| NeuroHarmonize - NeuroComBat | 0.06 [-0.7, 0.81] | 1.000 | 0.04 [-0.45, 0.52] | 1.000 |
| OPNested ComBat - NeuroComBat | 0.26 [-0.5, 1.01] | 0.970 | -0.03 [-0.52, 0.45] | 1.000 |
| AutoComBat - NeuroComBat | 0.39 [-0.36, 1.15] | 0.762 | -0.14 [-0.63, 0.34] | 0.988 |
| RELIEF - NeuroComBat | -0.25 [-1, 0.51] | 0.976 | 0.24 [-0.24, 0.73] | 0.803 |
| CovBat - NeuroComBat (all features) | -4.86 [-5.61, -4.1] | < 0.001 | 0.73 [0.24, 1.21] | < 0.001 |
| NeuroHarmonize - NeuroComBatAllFeatures | -5.25 [-6.01, -4.5] | < 0.001 | 0.9 [0.42, 1.39] | < 0.001 |
| OPNested ComBat - NeuroComBat (all features) | -5.05 [-5.81, -4.3] | < 0.001 | 0.83 [0.35, 1.32] | < 0.001 |
| AutoComBat - NeuroComBat (all features) | -4.92 [-5.67, -4.16] | < 0.001 | 0.73 [0.24, 1.21] | < 0.001 |
| RELIEF- NeuroComBat (all features) | -5.56 [-6.31, -4.8] | < 0.001 | 1.11 [0.62, 1.6] | < 0.001 |
| NeuroHarmonize - CovBat | -0.4 [-1.15, 0.35] | 0.748 | 0.18 [-0.31, 0.66] | 0.958 |
| OPNested ComBat - CovBat | -0.2 [-0.95, 0.55] | 0.993 | 0.11 [-0.38, 0.59] | 0.998 |
| AutoComBat - CovBat | -0.06 [-0.82, 0.69] | 1.000 | 0 .01[-0.49, 0.48] | 1.000 |
| RELIEF - CovBat | -0.7 [-1.46, 0.05] | 0.090 | 0.38 [-0.1, 0.87] | 0.251 |
| OPNested ComBat - NeuroHarmonize | 0.2 [-0.55, 0.95] | 0.993 | -0.07 [-0.55, 0.42] | 1.000 |
| AutoComBat - NeuroHarmonize | 0.34 [-0.42, 1.09] | 0.877 | -0.18 [-0.66, 0.31] | 0.956 |
| RELIEF - NeuroHarmonize | -0.3 [-1.06, 0.45] | 0.927 | 0.21 [-0.28, 0.69] | 0.905 |
| AutoComBat - OPNested ComBat | 0.14 [-0.62, 0.89] | 0.999 | -0.11 [-0.59, 0.38] | 0.998 |
| RELIEF - OPNested ComBat | -0.5 [-1.26, 0.25] | 0.468 | 0.27 [-0.21, 0.76] | 0.677 |
| RELIEF - AutoComBat | -0.64 [-1.39, 0.11] | 0.166 | 0.38 [-0.1, 0.87] | 0.246 |

**Supplementary Table 6:** Uncorrected BAG and MAE per harmonisation method for validation and testing sets, obtained using ASL-only or T1w+FLAIR+ASL features to predict brain age. Additional results of harmonisation of all features using NeuroComBat have been included under ‘NeuroComBat (all features)’. Note that the validation set was not corrected for age-bias. *ASL: arterial spin labelling; BAG: brain-predicted age gap; FLAIR: fluid attenuated inversion recovery; MAE: mean absolute error; T1w: T1-weighted.*

| **ASL-only** | **BAG**  **(μ ± σ)** | **MAE**  **(μ ± σ)** | **R^2^** |
| --- | --- | --- | --- |
| Unharmonised | -10.46 ± 14.72 | 13.24 ± 12.28 | 0.02 |
| NeuroComBat | -13.73 ± 10.21 | 14.78 ± 8.62 | 0.05 |
| CovBat | -13.82 ± 10.19 | 14.86 ± 8.61 | 0.05 |
| NeuroHarmonize | -13.92 ± 10.3 | 15.08 ± 8.52 | 0.03 |
| OPNested ComBat | -13.44 ± 10.46 | 14.66 ± 8.68 | 0.04 |
| AutoComBat | -12.02 ± 9.96 | 13.31 ± 8.17 | 0.08 |
| RELIEF | -16.01 ± 11.63 | 17.01 ± 10.12 | 0.02 |
| **T1w+FLAIR+ASL** | **BAG**  **(μ ± σ)** | **MAE**  **(μ ± σ)** | **R^2^** |
| Unharmonised | -4.70 ± 7.88 | 7.40 ± 5.42 | 0.30 |
| NeuroComBat | -4.70 ± 8.21 | 7.49 ± 5.78 | 0.31 |
| CovBat | -4.38 ± 8.14 | 7.28 ± 5.70 | 0.32 |
| NeuroHarmonize | -4.71 ± 8.24 | 7.51 ± 5.81 | 0.31 |
| OPNested ComBat | -4.50 ± 8.23 | 7.37 ± 5.80 | 0.31 |
| AutoComBat | -4.45 ± 8.13 | 7.31 ± 5.70 | 0.32 |
| RELIEF | -5.01 ± 8.45 | 7.74 ± 6.05 | 0.30 |
| NeuroComBat (all features) | -0.18 ± 6.20 | 4.71 ± 4.02 | 0.49 |
